# Generative design of antibody Fc interactions by deep mutational learning

**DOI:** 10.1101/2025.10.10.681689

**Authors:** Edward B. Irvine, Evangelos Stamkopoulos, Thomas Bikias, Joshua Kämpf, Lester Frei, Nick Schürmann, Albin Thaqi, Helen Schmid, Dimitri Coukos, Huilin Yang, Neda Nedeva, Raphael Pauli, Jonas Blatter, Chiara Rodella, Annmaree K. Warrender, Mason Minot, Giuliano Berloffa, Derek M. Mason, William Kelton, Sai T. Reddy

**Author notes:** Correspondence (EBI), (WK), (STR). Equal contribution (Edward B. Irvine, Evangelos Stamkopoulos, Thomas Bikias).

## Abstract

Antibodies have transformed modern medicine. Their remarkable therapeutic success stems from the fact that they do two things at once. The variable (Fab) domain recognizes a target, while the constant (Fc) domain determines how that recognition event is acted on by the immune system. The Fc carries out these instructions by interacting with a complex network of Fc-receptors expressed across diverse immune cells. Yet the ability to rewrite these instructions remains limited, as Fc-receptors bind overlapping surfaces on the Fc, making it difficult to engineer antibodies with precise patterns of receptor engagement. Here we introduce a new paradigm for Fc engineering, combining high-throughput experimental screening with deep learning to design antibodies with programmable patterns of Fc-receptor engagement. We constructed a library of more than 400 million Fc-variants and screened it against eight Fc-receptors by yeast display and deep sequencing, generating a high-resolution map linking Fc sequence to patterns of receptor recognition. Using these data, we developed FcGPT, a generative protein language model that designs millions of synthetic Fc-variants with user-defined Fc-receptor binding “profiles”. Experimental testing of 2,000 designs across 19 objectives yielded Fc-variants matching prescribed profiles of binding and non-binding across all eight receptors, including synthetic profiles not represented in nature. Success rates ranged from 5% to 100% among successful design tasks, including variants that selectively engage FcγR3A while avoiding FcγR3B (and vice versa), despite their >95% sequence identity. This work extends protein design from creating binders to individual targets to programming precise patterns of molecular recognition across entire families of receptors, offering a new way to engineer interactions between proteins and the immune system.

## INTRODUCTION

Monoclonal antibodies have profoundly transformed modern medicine^1^. Their success stems not only from their ability to bind specific antigens with high affinity, but also from their capacity to drive immune functions via their constant (Fc) domains. Through engagement with Fc-receptors, the antibody Fc domain orchestrates a wide range of immune functions that broadly serve to enhance pathogen clearance, modulate inflammation, and restore immune balance^2–4^. There are four natural human IgG Fc subclasses (IgG1, IgG2, IgG3, and IgG4), with IgG1 being the predominant choice for therapeutic use due to its robust effector function and stability^5^. While these subclasses differ in their natural effector profiles, Fc engineering enables the modulation of immune activity beyond what is afforded by subclass or allotypic variation alone^2^. Indeed, modifying the Fc domain has made a significant clinical impact, with Fc-engineered antibodies demonstrating improved therapeutic efficacy in areas such as oncology, autoimmunity, and infectious diseases^6–9^. Among currently marketed or late-stage clinical antibodies containing an Fc domain, nearly half incorporate Fc modifications^10^, reflecting a growing interest in harnessing Fc engineering for therapeutic benefit.

Fc engineering campaigns have traditionally focused on altering Fc-receptor binding affinity or specificity via glycoengineering, structure-guided mutagenesis, or directed evolution^7,11–17^. While these approaches have yielded effective variants, they remain limited in their ability to independently modulate binding across multiple Fc-receptors. This is in part due to the biological constraints of the system – many Fc-receptors bind to overlapping regions, so even single mutations often influence binding to multiple receptors simultaneously^2,18^. As a result, previous efforts offered limited control over the specificity, balance, and breadth of the Fc-mediated immune response. This limitation is clearly reflected in the clinic. Beyond antibodies designed to silence Fc function, nearly all Fc-engineered antibodies either: (i) enhance antibody-dependent cellular cytotoxicity (ADCC), or (ii) modulate antibody stability or half-life^19^.

Through engagement with Fc-receptors, the Fc domain can direct a far broader range of immune activities, including phagocytosis, cytokine release, antigen presentation, and immune cell polarization – each of which could be precisely tuned to combat disease^2–4^. Furthermore, consider that IgG antibodies interact with at least nine distinct Fc-receptors^3,4^. In a simplified model in which an antibody Fc can exhibit one of three binding states (e.g., enhanced, wild type, or no binding) to each receptor, this yields 3^9^, or nearly 20,000 unique Fc-receptor binding profiles, each potentially representing a unique way to tune immune function.

Unlocking this vast and largely unexplored functional space requires a detailed understanding of how Fc sequence encodes receptor engagement and downstream function. Experimental Fc engineering approaches are limited by the number of variants that can be screened^11–16^, making it intractable to evaluate the functional impact of combinatorial mutations across a domain as large as the antibody Fc. Furthermore, these approaches are inherently anecdotal – they focus on finding individual variants rather than uncovering the general rules that govern Fc sequence-function relationships.

Deep learning offers a powerful way to overcome these challenges. By uncovering latent, high-dimensional patterns from complex biological data, it can provide deeper insight into sequence-function relationships and guide advanced protein engineering efforts^20,21^. For instance, deep mutational learning, which combines high-throughput experimental screening with deep learning, enables the prediction of protein binding from sequence, including for previously untested combinations of mutations^22–26^. Beyond classification, recent advancements in natural language processing^27–29^, have inspired the development of protein language models (PLMs) for sequence representation and generation^21,30–33^. PLMs model the biochemical “language” of proteins by learning residue distributions through autoregressive or masked pre-training on vast protein sequence datasets. This training process captures the thermodynamic and evolutionary constraints embedded in protein sequences, enabling the computational design of proteins with a high likelihood of proper folding and function^31,34^. Notably, while such classification and generative strategies are increasingly applied to the antibody variable domain to optimize antigen binding ^35^, they have not yet been applied to the antibody constant region.

In this study, we establish a new approach for Fc engineering that combines high-throughout experimental screening with generative artificial intelligence (**Extended Data Fig. 1**). We performed deep mutational scanning (DMS) across the entire 232 residues of the IgG1 Fc domain to determine how single mutations impact binding to eight canonical Fc-receptors. These data guided the design of an Fc combinatorial library of more than 400 million variants, with mutations focused on regions critical for Fc-receptor engagement. We then screened this library for binding to eight Fc-receptors, creating a large-scale atlas of Fc:Fc-receptor binding interactions which was used to train deep learning classifiers that accurately predict Fc-receptor binding based on sequence. To move from prediction to design, we developed FcGPT, a generative protein language model trained on our Fc dataset and optimized through reinforcement learning^36–38^. Experimental validation of 2,000 designs across 19 receptor interaction profiles demonstrated that FcGPT can generate novel Fc-variants with prescribed binding and non-binding across eight Fc-receptors simultaneously. These include synthetic profiles not represented by natural IgG subclasses that have been difficult to create through conventional engineering approaches. Beyond Fc biology, this work extends protein science from designing proteins that bind individual targets, to designing interactions across entire networks of receptors.

## RESULTS

### Establishing an antibody Fc surface display system

To enable screening of antibody Fc-variant libraries, we utilized an antibody Fc surface display system in yeast. Antibodies are modular molecules comprised of a Fab domain, which regulates antigen binding, and an Fc domain, which interacts with Fc-receptors to trigger effector functions. Previous studies have shown that isolated Fc domains remain functional even in the absence of a Fab domain^39–42^. This functionality is retained in yeast display systems^39^, which allow for the efficient coupling of genotype and phenotype at high-throughput^43,44^. Here, our yeast system encodes a copy of the Aga1 protein linked via disulfide bonds to a vector-encoded copy of the Aga2 protein fused to the human IgG1 Fc domain (**Fig. 1a**). HA and FLAG tags were additionally included to enable detection of surface display by flow cytometry (**Fig. 1a**).

**Fig. 1.**
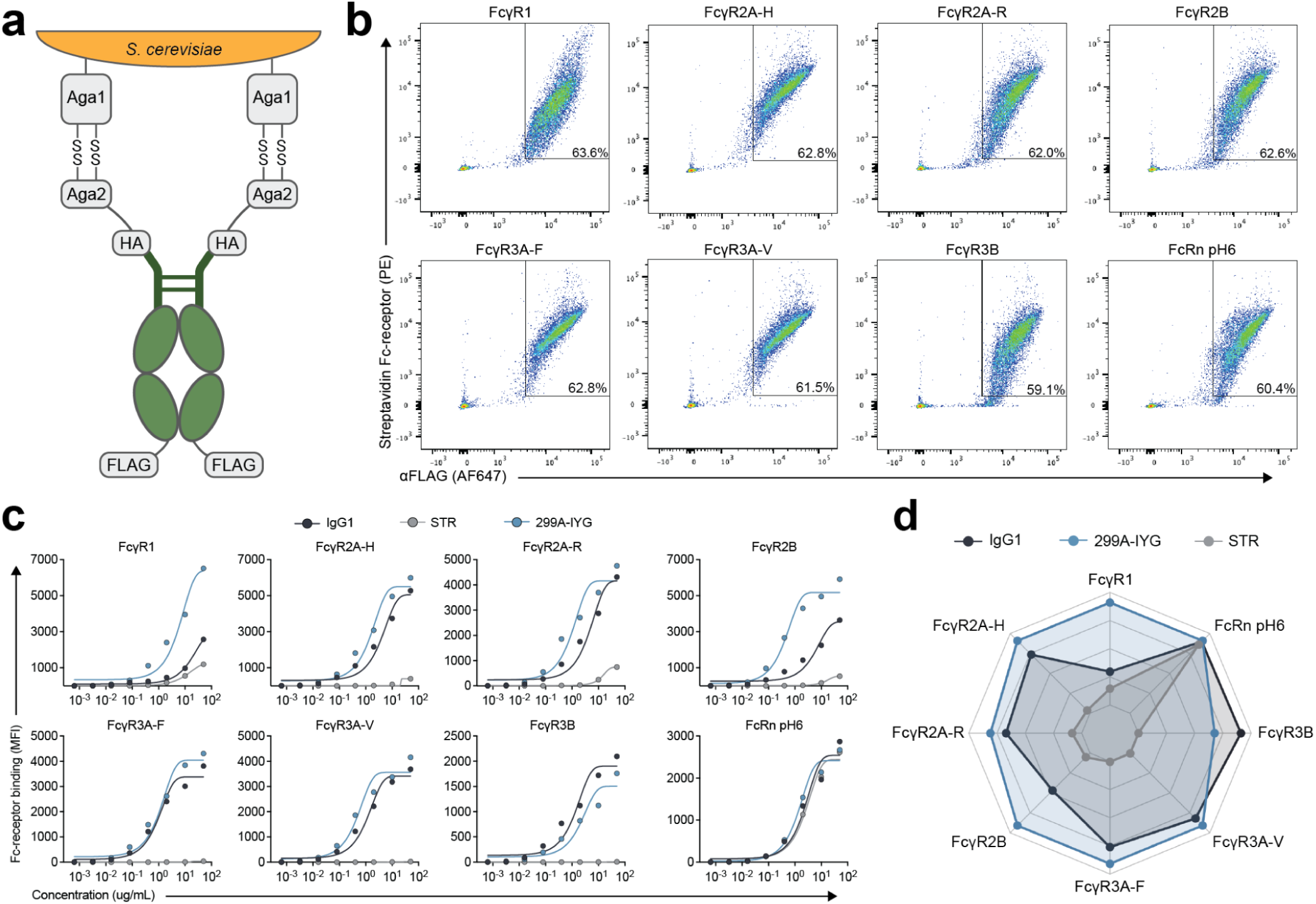
Development mid characterization of an antibody Fc surface display sy stem, a, Schematic of lite antibody Fc yeast surface display approach. Image created with BioReruter.com. b. Flow cytometry analysts of Fc-receptor binding in the yeast display system for wild-type human IgG I Fc. X-axis: FLAG surface expression. Y-axis: Fc-receptor binding. Fc-receptors were used at a concentration of 5Dpg’ntl.. FcRn staining was performed al pH6. c, IgGL aglycosylated Fc-variant 299A-IYG, and Fc null variant STR were stained across a titration series of each Fc-receptor. X-axis: Fc-receptor concentration. Y-axis: Fc-receptor binding (geometric mean fluorescence intensity) normalized to FLAG surface expression. Data were fitted using a sigmoidal (4-parameter logistic) curve, d, Radar plot showing the area under the binding curves from the Fc-receptor titration in panel c for IgGL 299A-1YG. and STR,

To assess expression and functionality, yeast displaying the human IgG1 Fc domain were stained with a panel of eight canonical Fc-receptors: FcγR1, FcγR2A-H (167H), FcγR2A-R (167R), FcγR2B, FcγR3A-F (176F), FcγR3A-V (176V), FcγR3B, and FcRn (**Extended Data Table 1**). Robust surface expression and Fc-receptor binding were observed across each of the Fc-receptors by flow cytometry (**Fig. 1b**), confirming that the displayed Fc domain retained structural integrity and binding activity. To determine the ability of the system to detect functionally relevant differences in Fc-receptor engagement, we next expressed the Fc-silencing, STR variant (L234S/L235T/G236R) on the yeast surface^45,46^. This variant is known to abrogate binding to all Fcγ receptors while retaining interaction with FcRn at pH of 6^45,46^. Yeast displaying the STR variant were stained with soluble Fc-receptors across a range of concentrations and compared to wild-type human IgG1 (**Fig. 1c**). As expected, STR showed robust FcRn binding but minimal interaction with Fcγ receptors (**Fig. 1c,d**), demonstrating that the Fc display system can resolve differences in Fc functionality.

Beyond Fc sequence, Fc glycosylation is also a critical determinant of antibody Fc function^2^. Of note, yeast express proteins with more mannose-rich glycosylation patterns than mammalian cells^47^, which could affect Fc-receptor binding and potentially confound interpretation of sequence-driven functional effects. To eliminate this variable, we also displayed the aglycosylated, 299A-IYG variant (T299A/K326I/A327Y/L328G) on the yeast surface^48^. While mutating the N-X-S/T motif required for Fc glycosylation typically reduces binding to the low affinity Fcγ receptors^49,50^, 299A-IYG was engineered to be aglycosylated while retaining near-wild-type IgG1 binding to the canonical Fcγ receptors^48^. Consistent with prior reports, 299A-IYG exhibited robust binding across all Fc-receptors tested (**Fig. 1c,d**)^48^. This glycan-independent functionality enables Fc:Fc-receptor binding interactions to be assessed in the absence of yeast-specific glycosylation. Hence, we selected 299A-IYG as the scaffold for our display system and used it in all subsequent experiments. Collectively, these data establish a robust protein display platform for high-throughput functional screening of antibody Fc-variants.

### Deep mutational scanning of the Fc domain

After establishing a platform for antibody Fc surface display, we next aimed to create a map of Fc sequence-function relationships to guide the design of a large Fc combinatorial library. Recent work profiled the effect of single amino acid substitutions in the human IgG1 domain on Fcγ receptor binding^42^. However, residues critical for Fc-receptor binding in an aglycosylated genetic background remain undefined. To systematically assess the functional impact of single-site mutations in the aglycosylated 299A-IYG background, we performed deep mutational scanning (DMS) by single-site tiling of degenerate codons (NNK, N=A,C,G,T; K = G,T) to allow substitutions with any of the 20 canonical amino acids across residues E216 to K447 (EU numbering) of the Fc domain (**Fig. 2a**). Only the alanine at position 299 was held constant to preserve the aglycosylated background.

**Fig. 2.**
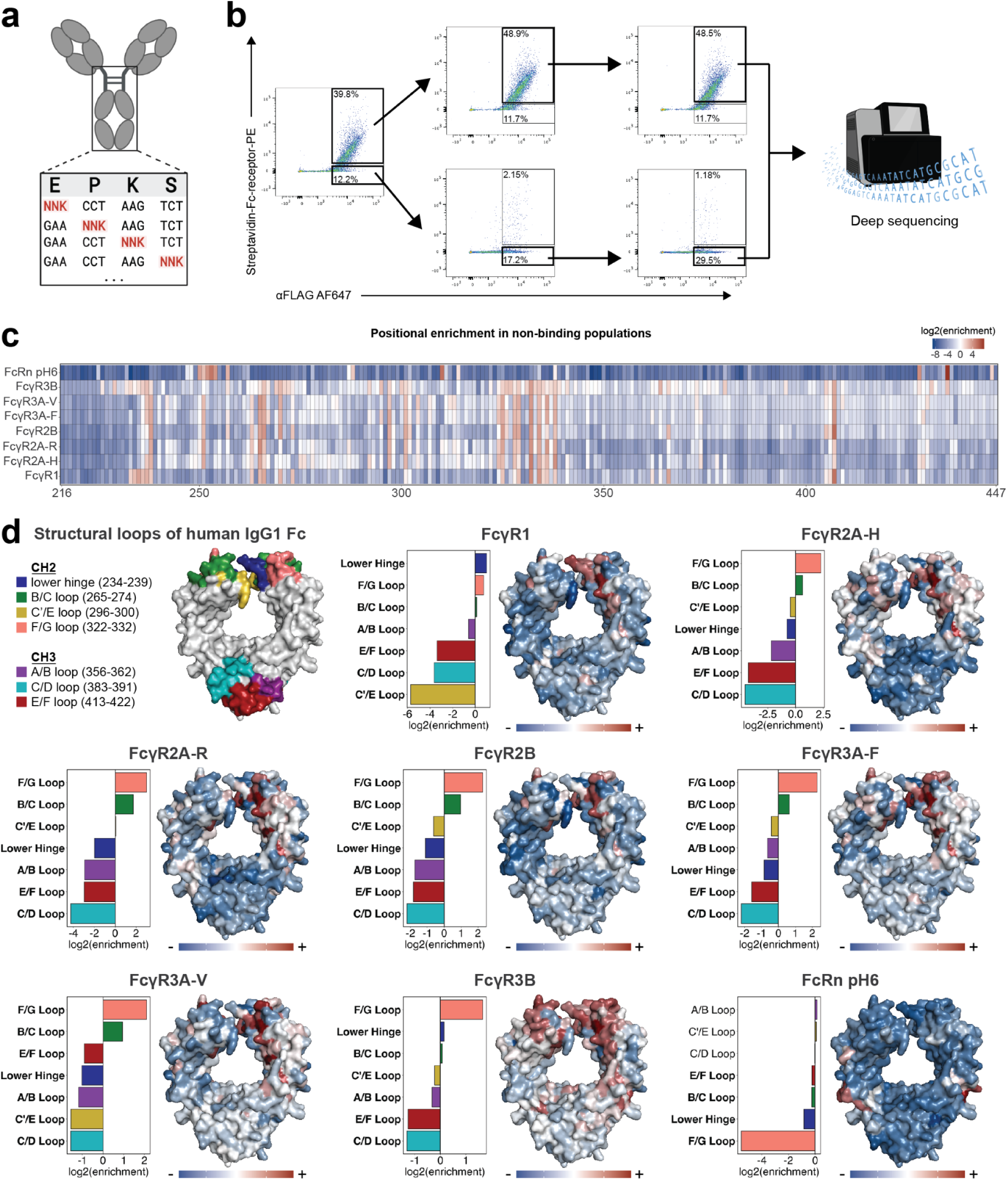
Deep mutatlnnal scanning of the Fc dumain. a, Schematic of smgle-site saturation mutagenesis across the Fc domain using NNK codons. b, FACS-based screening workflow (FcγR3A-F shown), with iterative sorting of binding and non-binding populations followed by deep sequencing, c, Heatmap showing Iog2 enrichment of mutations in non-binding populations relative to the naive library across eight Fc-receptors. Columns indicate Fc positions (EL^1^ numbering), positive enrichment identifies residues important for receptor binding, d. Structural mapping of positional enrichment onto human lgG1 Fc (PDB: 3AVE). Top left, Fc structure highlighting known structural loops. Remaining panels show positional enrichment (red, positive; blue, negative) and regional enrichment of structural loops (bar plots) Hold labels show significant enrichment (Fisher s exact test, Bonfemxii’adjusied P*-0.05). Panel a and pan of panel b created with BioRender corn

Deep sequencing of the DMS library revealed approximately 90% coverage of all possible single-site amino acid mutations (**Extended Data Fig. 2a**). We then screened our yeast Fc DMS library for binding across a panel of eight Fc-receptors via fluorescence-activated cell sorting (FACS), isolating Fc-receptor binding and non-binding populations through multiple rounds of selection (**Fig. 2b**). The sorted populations were deep sequenced to quantify Fc-variant frequency in the binding and non-binding populations compared to the naïve (unsorted) library (**Fig. 2c** and **Extended Data Fig. 2b**). Analysis of the non-binding populations, which are depleted of Fc-variants that retain receptor engagement, highlighted residues essential for Fc-receptor binding. Variants enriched in this population are those that disrupt receptor interaction, allowing positions critical for Fc:Fc-receptor interaction to be identified. We observed strong enrichment at multiple control positions, validating the fidelity of the screen. For example, L234 and L235 were positively enriched in the non-binding population across multiple Fcγ receptors (FcγR1, FcγR2A-H, FcγR2B, FcγR3A-F, FcγR3A-V, FcγR3B) (**Fig. 2c**), consistent with established Fc silencing variants such as STR (L234S/L235T/G236R) and LALA (L234A/L235A), which carry mutations at these positions^45,51^. Similarly, D265, a residue known to be critical for Fcγ receptor engagement^18^, was consistently enriched in the non-binding populations across all Fcγ receptors tested (**Fig. 2c**), reinforcing its functional importance. Finally, H310 and H435, critical determinants of pH-dependent FcRn binding^52^, were strongly enriched in the FcRn non-binding population (**Fig. 2c**), confirming their importance in the aglycosylated 299A-IYG context as well.

Beyond canonical receptor-contact sites, our screen identified additional sites critical for Fcγ receptor engagement outside of well-established residues. For instance, L251 was positively enriched in the non-binding populations for FcγR2A-H, FcγR2A-R, FcγR2B, FcγR3A-F, and FcγR3A-V (**Fig. 2c**), suggesting its importance for interacting with these receptors. Similarly, residues K334, I336, and K338 were consistently enriched in the non-binding populations across all Fcγ receptors tested (**Fig. 2c**), pointing to their importance in Fc:Fcγ receptor interactions in the aglycosylated 299A-IYG background.

To contextualize these data, we mapped enrichment scores from the non-binding populations onto a high-resolution human IgG1 Fc crystal structure (**Fig. 2d**). In the context of human IgG1, the lower hinge and three structural loops in the CH2 domain – the B/C, C′/E, and F/G loops – are known to contain residues that directly contact Fcγ receptors^18,48,53,54^. Consistent with previous work, we found the F/G loop to be significantly positively enriched for mutations across all Fcγ receptors (**Fig. 2d**), supporting its critical role in Fcγ receptor recognition in the 299A-IYG background. The B/C loop was also consistently significantly positively enriched, albeit with smaller effect sizes (**Fig. 2d**). The lower hinge region was significantly positively enriched in the FcγR1 and FcγR3B datasets, suggesting that an extended portion of this region contributes to binding these receptors (**Fig. 2d**). By contrast, CH3-domain loops were significantly negatively enriched across all Fc-receptors as expected (**Fig. 2d**), suggesting their limited involvement in Fc-receptor binding in this aglycosylated context. Together, these data reveal how single-site mutations across the aglycosylated 299A-IYG Fc domain influence binding to eight canonical Fc receptors, identifying contributions from both previously characterized residues and previously unrecognized positions.

### Design and construction of a 400-million-variant Fc library

To explore how combinations of mutations influence Fc-receptor binding, we used our DMS data to rationally design a combinatorial Fc library. We defined “high-leverage” positions as those exhibiting significant enrichment in the non-binding population for at least one Fc receptor and selectively mutated these positions (**Fig. 3a**). Conversely, “low-leverage” residues which were not significantly positively enriched, were left as wild type. We performed this segregation of residues independently for each Fc-receptor, resulting in a map of residue importance spanning residue P232 – L368 (EU numbering) of the Fc domain for each Fc-receptor evaluated (**Fig. 3a**). Residues V369 – K447 had a limited number of significantly positively enriched positions (**Fig. 2c**), so this region was left unmutated enabling Illumina-based deep sequencing of the library.

**Fig. 3.**
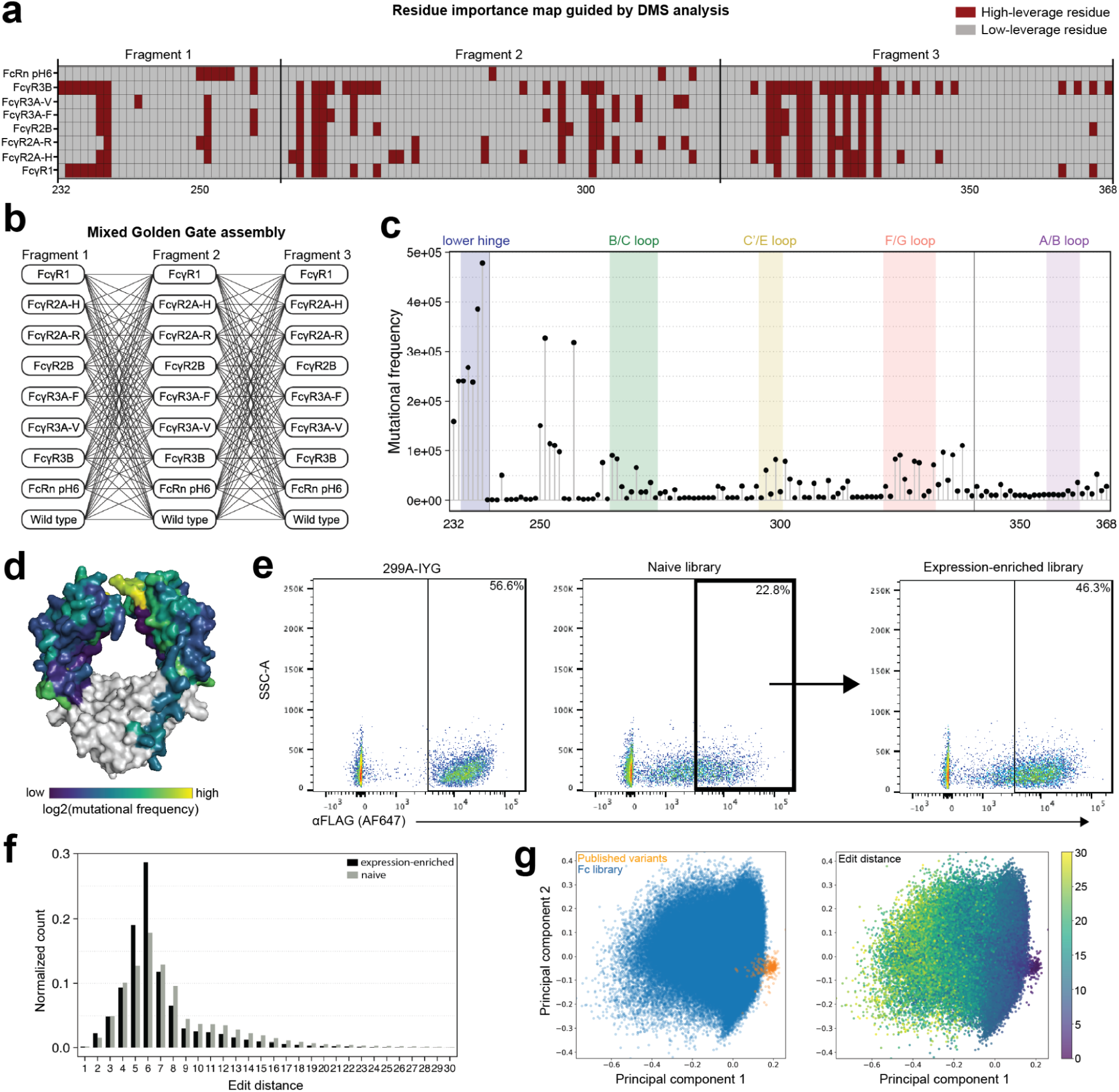
Rational design and cniwtniction of a diverse Fc combinatorial library, a, DMS-guided map of Fc residue importance across eight receptors (EU numbering). High-leverage residues (red) were identified by significant enrichment of mutations in non-binding populations (Fisher’s exact test, Bonferroni-adjusted P<0.05); low-leverage residues arc shown in grey, b, Mixed Golden Gate assembly of three fragment pools, each containing 70% wild-type and 30% distributed across eight receptor-guided mutational variants. C, Mutational frequency across Fc positions (EU numbering), with structural loops highlighted d. Structural mapping of log: mutational frequency onto human IgGI Fc (PDU 3AVE). e. Flow cytometry showing FLAG surface expression in the naive and expression-enriched libraries, with parental 299A-IYG shown for reference f, Edit distance front 299A-IYG in naive (grey) and expression-enriched (black) libraries g, PC A of ESM2 sequence embeddings complin ng the exprcssion-ennchcd Fc library with JOO previously published Fc-vanants Points are colored by source (left) or edit distance from 299A-IYG (right).

To bias mutagenesis towards variants likely to retain or enhance receptor binding, we used custom degenerate codons at high-leverage sites reflecting the amino acid distributions observed in the Fc-receptor binding populations from DMS, rather than standard NNK codons (**Extended Data Fig. 3a**). Based on this design, we generated 24 unique, mutant library fragments *in silico* (three designed based on the DMS data from each Fc-receptor), and ordered them as single-stranded oligonucleotides. To accommodate the length constraints of the oligonucleotides, the library was divided into three fragments. Together, these design choices yielded a combinatorial library with a theoretical sequence space of 10^62^ variants.

To assemble the full-length Fc combinatorial library, we combined the 24 mutational fragments with their corresponding wild type (299A-IYG) counterparts in a single Golden Gate reaction^55^ (**Fig. 3b**). Each of the three Fc domain fragments thus had nine potential states: eight receptor-specific mutant versions and one wild type version (**Fig. 3b**). This approach allowed us to carefully control library composition by simply altering the ratio of mutant to wild type fragments. Ultimately, to maintain a controlled edit distance from wild type, we included 70% of the wild type fragments in the Golden Gate assembly reaction, distributing the remaining 30% evenly among the eight mutational fragments.

The resulting library was then co-transformed with a linearized pYD1 vector into yeast for assembly via homologous recombination, yielding 4.1×10^8^ transformants. Deep sequencing revealed that mutations were commonly located in the lower hinge and CH2 domain loops (B/C, C′/E, and F/G) (**Fig. 3c,d**). As expected, strong and significant correlations were observed between the location of mutations in the library and the expected location of mutations based on the library design (**Extended Data Fig. 3b**). Although a modest bias towards mutations in fragment 1 was observed, leading to a slight overrepresentation of mutations at the N-terminus of the library (**Fig. 3c** and **Extended Data Fig. 3b**).

Flow cytometry analysis of the yeast Fc combinatorial library revealed that approximately 23% of the library exhibited surface expression, compared to 57% for the 299A-IYG variant (**Fig. 3e**). To enrich for well-folded, surface-expressing Fc-variants, we sorted the library for expression, isolating an expression-enriched population for downstream analyses (**Fig. 3e**). This enriched library had a median edit distance of six from wild type, one less than the naive library (**Fig. 3f**).

To contextualize the diversity of our Fc combinatorial library, we compiled a comprehensive dataset of 301 unique human IgG1 Fc-variant sequences from the published literature (**Supplementary Table 1**). These variants, along with the sequences comprising our Fc combinatorial library, were embedded using the ESM2 protein language model^32^, and visualized by uniform manifold approximation and projection (UMAP) (**Fig. 3g**). Notably, our library spans a substantially broader and more diverse region of Fc sequence space than the set of previously reported variants (**Fig. 3g**), providing a unique resource for discovering Fc-variants with novel functionalities.

### Deep learning models accurately predict Fc-receptor binding based on sequence

Having constructed a large and diverse Fc combinatorial library enriched for surface expression, we next sought to develop classifiers capable of linking Fc sequence with Fc-receptor binding activity (**Fig. 4a**). To this end, we systematically screened the Fc combinatorial library across our panel of eight canonical Fc-receptors, isolating binding and non-binding populations for each via FACS (**Extended Data Fig. 4a**). To further enrich for variants with selective Fc-receptor binding profiles, we performed an additional FACS campaign, in which the library was stained simultaneously with pairs of Fc-receptors, enabling the direct isolation of variants that bound one receptor but not the other (**Extended Data Fig. 4b**). This approach encompassed all six pairwise combinations of FcγR1, FcγR2A-R, FcγR2B, and FcγR3A-F. Sorted populations were then deep sequenced, filtered to remove low-quality sequences and variants supported by fewer than three sequencing reads, and translated into amino acid sequences. Combining data from these combinatorial library screens with the Fc DMS dataset described above yielded a training dataset comprising 1,474,696 unique Fc sequences (DMS - 324,092; combinatorial - 1,101,665; combinatorial co-stain - 48,939).

**Fig. 4.**
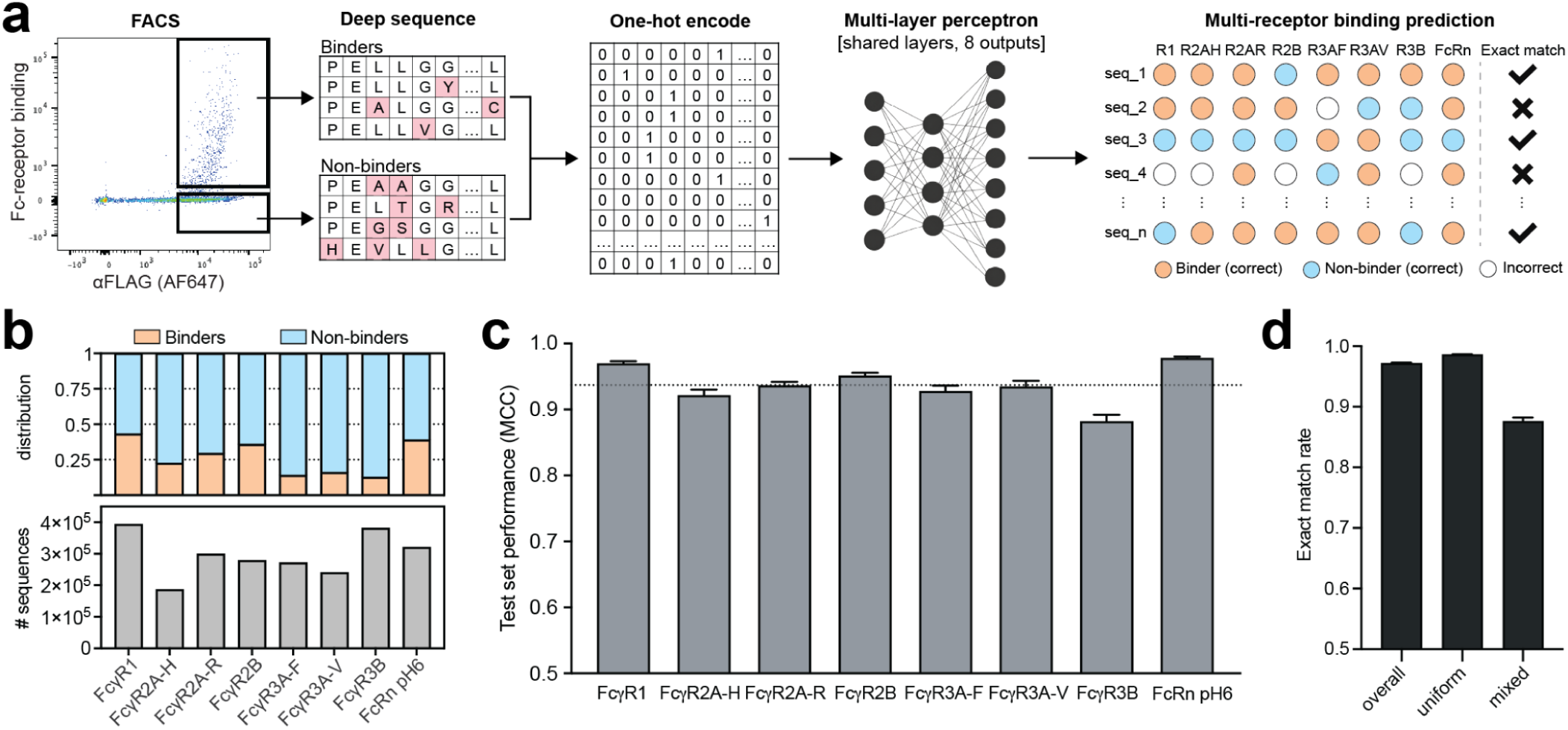
Integrating FACS and deep sequencing with machine learning to predict Fc-reeeptnr binding, a. Binding and non-bindmg populations for each Fc-receptor were isolated via FACS and deep sequenced. Fc sequences were one-hot encoded and used to train a multilayer perceptron (MLP) with shared hidden layers and eight receptor-specific outputs to predict binding across the Fc-receptor panel b, Number (bottom) and class distribution (top) of training data following preprocessing, c, Ensemble classifier performance on held-out test data per Fc-receptor shown by Matthew’s Correlation Coefficient (MCC) Dotted line indicates the mean MCC across all eight receptors (0.9)7) d. Exact match rate, defined as the proportion of test sequences for which all experimentally determined Fc-receptor labels were predicted correctly. Results are shown for the overall test set. as well as for Fc sequences with uniform (all the same) or mixed (at least one different) binding labels across receptors.

For the six low affinity Fcγ receptors, the resulting data were dominated by non-binding Fc sequences, whereas those for FcγR1 and FcRn pH6 showed a more balanced distribution of binders and non-binders (**Fig. 4b**). As expected, given that the throughput of FACS (∼10^7^ cells) undersamples the full library (∼10^8^ variants), the majority of sequences had experimental binding data for only one Fc-receptor (**Extended Data Fig. 4c**). Nevertheless, many Fc sequences contained binding data across multiple Fc-receptors, with 5,066 fully labeled across all eight (**Extended Data Fig. 4c**).

After compiling the full dataset of 1.5×10^6^ unique Fc-variants, the sequences were encoded and used to train machine learning models that predict binding across the eight Fc-receptors. For initial benchmarking, we systematically evaluated multiple model architectures, including logistic regression, multilayer perceptrons (MLPs), convolutional neural networks (CNNs), and transformer-based models, using sequence representations ranging from simple one-hot encoding to embeddings from the pre-trained protein language model ESM-C^56^ (**Extended Data Table 2**). Hyperparameters were optimized using the tree-structured Parzen estimator algorithm^57^. Models were trained on 85% of the dataset, validated on 10%, and evaluated on the remaining 5% held out for testing. Because a random split can reward memorization of shared mutations, we also tested three leakage-safe splits that hold out whole mutation combinations, clusters of spatially adjacent mutated positions, or the most frequently mutated positions. Model performance was evaluated using Matthew’s correlation coefficient (MCC) to account for imbalances in class size^58^. Among the architectures evaluated, a multi-layer perceptron (MLP) trained on one-hot-encoded Fc sequences achieved the highest predictive performance, outperforming logistic regression, CNNs, and transformer-based models (**Extended Data Table 2**). Trained on one-hot-encoded Fc sequences, this model also matched or exceeded the performance of classifiers using pretrained ESM-C embeddings across most data splits (**Extended Data Table 2**).

On the held-out test set, this classifier achieved an average Matthews correlation coefficient (MCC) of 0.937, with individual MCC values exceeding 0.88 across all eight Fc-receptors (**Fig. 4c**). To evaluate predictive performance across the entire panel of Fc-receptors, we next assessed the exact match rate, which we define as the proportion of sequences for which all experimentally determined Fc-receptor labels were predicted correctly. Exact match provides a stringent measure of model performance, as it requires the models to make correct predictions for every available Fc-receptor label for a given Fc sequence. Across the entire held-out test set, the classifier achieved an exact match rate of 0.972 (**Fig. 4d**). To further dissect performance in a multi-receptor context, we stratified the test set into sequences with uniform labels (all labeled Fc-receptors either bind or do not bind) and those with mixed labels (a combination of binding and non-binding Fc-receptor labels). While the exact match rate was highest for uniform-label sequences (0.986), performance remained strong even for the more challenging mixed-label group (0.876) (**Fig. 4d**), underscoring the ability of the models to robustly classify Fc-variants with complex, multi-receptor binding profiles. Taken together, these results demonstrate that deep learning models trained on high-throughput protein display data can decode the underlying Fc sequence-function relationships to accurately predict how even a small number of amino acid substitutions alter Fc-receptor binding, providing a powerful tool for predicting and ultimately engineering antibody Fc function.

### FcGPT designs Fc-variants with tailored Fc-receptor binding profiles

Given the ability of deep learning models to learn the hidden relationships between Fc sequence and function to accurately predict Fc-receptor binding (**Fig. 4c,d**), we hypothesized that this process could be inverted – that starting from a desired Fc-receptor binding profile, it should also be possible to design Fc-variants that match the target profile. However, because our Fc combinatorial library spans a theoretical sequence space of 10^62^ unique Fc-variants, identifying Fc-variants with rare binding profiles through simple *in silico* search is computationally prohibitive. We reasoned that a deep generative model would provide a more efficient way to explore this vast sequence space and design Fc-variants with the desired properties.

Inspired by recent advances in protein language models, which learn the biochemical grammar and syntax of proteins from large sequence corpora^21,30–33^, we sought to develop a domain-specific protein language model for computational Fc design. Protein language models have demonstrated a strong capacity for realistic *de novo* sequence generation, and conditional protein language models extend this ability by guiding sequence generation toward specified protein families or functional outcomes^30,31^. Building on this framework, we created FcGPT, a conditional protein language model trained to learn the biochemical grammar of the antibody Fc domain (**Extended Data Fig. 5a**).

FcGPT was pre-trained directly on our Fc dataset (1.5×10^6^ unique Fc-variants) using an autoregressive next-token prediction objective (**Extended Data Fig 5a**). The backbone of the model is a multi-layered GPT-J-style transformer decoder architecture where each layer includes a multi-head self-attention module and standard feed-forward neural networks preceded by layer normalization. During pre-training, we applied causal (left-to-right) masking for autoregressive sequence generation and rotary positional embeddings for improved relative position encoding. Analogous to ProGen2 where control tags are linked to protein sequences to enable conditional generation^30^, FcGPT prepends a fixed eight-token conditioning prompt to each Fc sequence (**Extended Data Fig 5a**). Each token corresponds to one Fc-receptor (FcγR1, FcγR2A-H, FcγR2A-R, FcγR2B, FcγR3A-F, FcγR3A-V, FcγR3B, and FcRn pH6) and encodes the experimentally determined binding status as binder (<yes>), non-binder (<no>), or unknown (<pad>) (**Extended Data Fig 5a**). Through this strategy, FcGPT simultaneously learns the amino acid distribution preset in our Fc library, and the underlying sequence dependencies associated with specific Fc-receptor binding profiles. Loss trajectories during model pre-training showed that larger models converged faster and achieved lower binary cross-entropy losses, but performance gains plateaued beyond ∼10 million parameters (**Extended Data Fig. 5b-d**). We therefore selected the 10 million parameter FcGPT model for all downstream tasks.

Pre-training enabled FcGPT to capture broad Fc sequence-function relationships. However, it was insufficient to reliably generate variants matching specific Fc-receptor binding profiles, likely due to label sparsity in the dataset – most Fc sequences only had binding data for a single Fc-receptor (**Extended Data Fig. 4c**). To address this, we utilized a reinforcement learning-based optimization strategy using experimental feedback (RLXF) (**Fig. 5a**)^38^. In this framework, FcGPT is treated as a policy model and optimized using Group Relative Policy Optimization (GRPO)^37^, the groupwise objective popularized by DeepSeek-R1 for scalable verifier-only reinforcement learning^36^. At each step, FcGPT generates batches of Fc-variants conditioned on a user-defined Fc-receptor binding profile (**Fig. 5a**). Analogous to the synthetic verifier in DeepSeek-R1, each proposed sequence is scored by our high-accuracy Fc-receptor binding MLP classifiers (**Fig. 4c,d**), which act as an experimentally trained, model-based verifier to produce scalar rewards for GRPO updates. The resulting predictions are used to construct a multi-objective reward function that: (i) rewards alignment with the specified receptor profile, (ii) penalizes excessive sequence divergence from the 299A-IYG wild type Fc, (iii) penalizes duplicate sequences, and (iv) penalizes mutations at positions the training library did not vary, focusing mutations where the binding classifiers are most reliable (**Fig. 5a**). GRPO further incorporates a KL-divergence regularization term between the fine-tuned policy and the pre-trained FcGPT, which stabilizes learning and prevents catastrophic drift from the base model distribution. To efficiently update parameters, we applied low-rank adaptation (LoRA) modules^59^, yielding lightweight Fc-profile-specific adapters while keeping the base FcGPT model frozen (**Fig. 5a**). Once aligned by RLXF, FcGPT could design millions of sequences for any target Fc-profile.

**Fig. 5.**
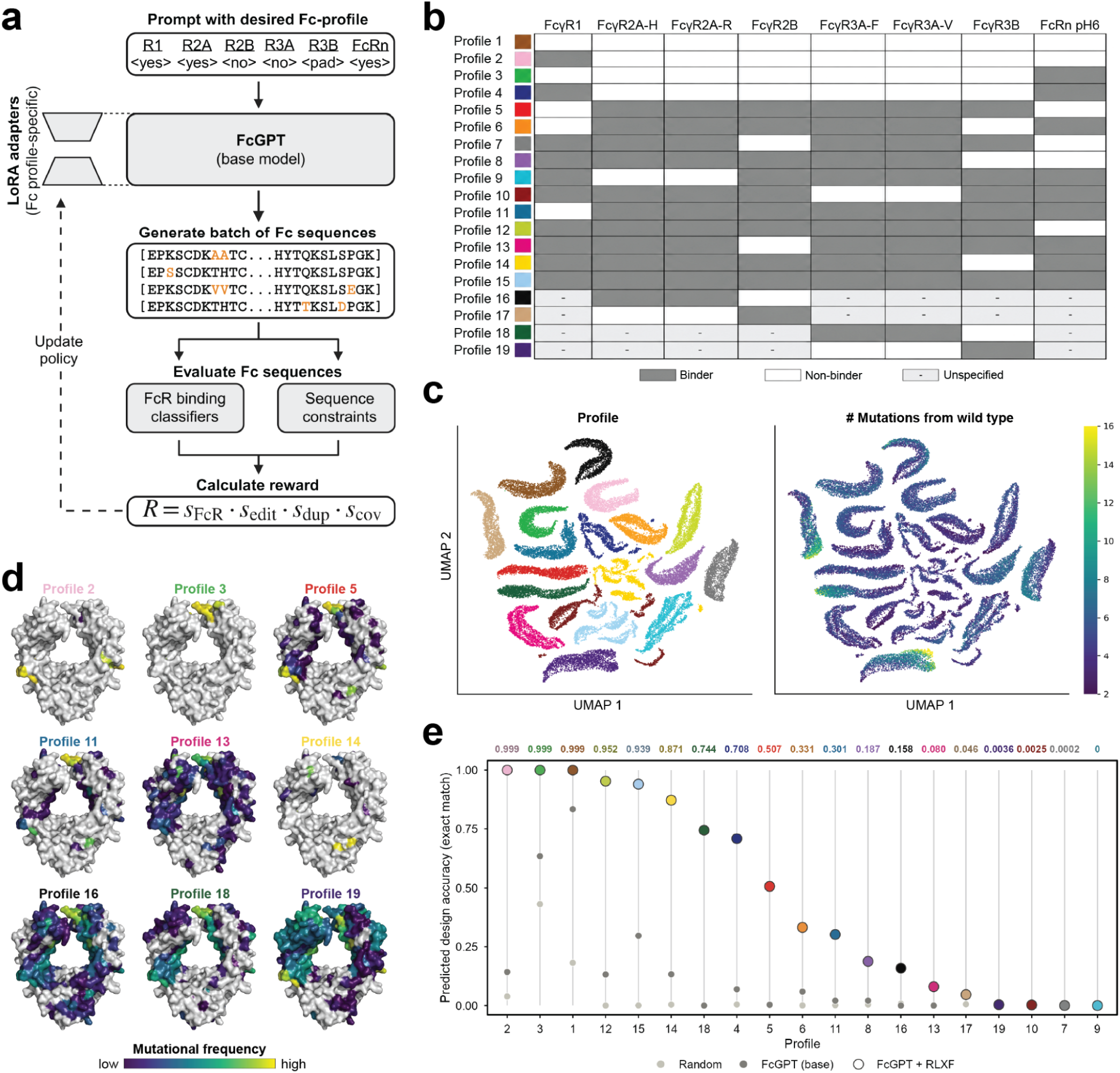
Reinforcement learning enables FcGPT to design antibodies with tailored Fc-receptor binding pruflles. a, Overview of reinforcement learning with experimental feedback (RLXF). FcGPT is prompted with a desired Fc-receptor binding profile and generates Fc sequences, which are evaluated using Fc-receptor binding classifiers and sequence constraints. A composite reward guides policy optimization using Group Relative Policy Optimization (GRPO) and profile-specific LoRA adapters, h, Nineteen target Fc-receptor binding profiles used for sequence design. Gray indicates binding, white indicates noo-binding, and dashes indicate unspecified binding preferences, c, UMAP projections of FcGPT-dtsigned sequences, colored by target binding profile (left) or number of mutations relative to wild-type IgGI (right), d, Mutational frequencies of designed sequences for representative profiles mapped onto the human IgGI Fc structure (PDB: JAVE). e, Predicted design accuracy across N target profiles fur random amino acid sampling, the FcGPT base model, and FcGPT optimized with RI.XF Accuracy is defined as the fraction of generated sequences matching all specified receptor-binding objectives. as evaluated by Fc-receptor binding classifiers Numbers above the plot indicate RIXF design accuracy for each profile,

To evaluate design performance, we tasked FcGPT with generating 2000 sequences across 19 diverse Fc-receptor binding profiles (**Fig. 5b**). These profiles spanned a broad range of functional objectives, from complete receptor silencing (Profile 1) to engagement of all eight Fc-receptors (Profile 15), as well as complex combinations of receptor binding and non-binding. We also included some of the most challenging design objectives in the field of Fc engineering, such as FcγR2A/FcγR2B-selective variants and FcγR3A/FcγR3B-selective variants, where these Fc-receptor pairs share >95% sequence identity in their extracellular domains^60,61^. Moreover, the majority of these profiles represent synthetic combinations of Fc-receptor binding and non-binding activities not found among natural human IgG subclasses, providing a stringent benchmark for the generative capabilities of FcGPT.

To visualize the distribution of the Fc-variants generated, we encoded the sequences using FcGPT-derived embeddings and projected them using UMAP (**Fig. 5c**). FcGPT-designed sequences clustered according to their target Fc-receptor binding profiles (**Fig. 5c**), indicating that distinct functional objectives guide the model towards different regions of Fc sequence space. Coloring sequences by their number of amino acid substitutions from wild type revealed that some profiles could be achieved with only a few substitutions (e.g., Profile 2), whereas others included designs with substantially greater sequence divergence (e.g., Profile 19) (**Fig. 5c**). Mapping the mutational frequency of FcGPT-generated variants onto the Fc crystal structure provided a structural view of the sequence changes underlying the different synthetic Fc-profiles (**Fig. 5d**). For example, variants designed to abrogate FcγR1 binding (e.g., Profiles 3, 5, and 11) showed a concentration of substitutions in the lower hinge (**Fig. 2d** and **5d**), consistent with established FcγR1 contact sites and our DMS screen (**Fig. 2d**), illustrating that FcGPT captures and reconstructs biologically meaningful sequence-function relationships during the design process.

Finally, we evaluated the design accuracy of FcGPT across all nineteen Fc-receptor binding profiles using the exact match rate predicted by our classifiers. We benchmarked FcGPT optimized through reinforcement learning (FcGPT-RLXF) against random sequence sampling and the pre-trained FcGPT-base model without reinforcement learning. Both benchmarks generally performed poorly, particularly for complex receptor-binding profiles, indicating that pre-training alone is insufficient for precise, targeted Fc-variant design (**Fig. 5e**). By contrast, FcGPT-RLXF generated sequences predicted to satisfy 18 of the 19 target profiles, achieving exact match rates exceeding 0.20 for eleven profiles, including eight exceeding 0.70 and three approaching perfect accuracy (0.999) (**Fig. 5e**). Notably, high design accuracy extended to receptor-selectivity objectives, including retention of FcγR3A binding while eliminating FcγR3B binding (e.g., Profile 14, 0.871). Accuracy was lower for other particularly challenging objectives, such as FcγR2A/FcγR2B-selectivity (e.g., Profiles 9 and 17). Encouragingly, these patterns mirror current experimental Fc engineering challenges, suggesting the model is capturing biologically meaningful constraints in the Fc sequence-function landscape. Together, these results establish FcGPT as a generative model that combines high-resolution protein mutagenesis data with deep learning to design antibodies with user-defined binding properties.

### Experimental validation of FcGPT-designed Fc-variants

We next sought to determine whether FcGPT could translate specified patterns of receptor binding and non-binding activity into Fc sequences that exhibited those properties in the lab. To test this, we used the RLXF-optimized FcGPT model to generate over 9 million candidate Fc sequences spanning the nineteen target profiles (**Fig. 5b**). We used two parallel design strategies, employing either the ensemble of Fc-receptor binding classifiers described above, or a separately trained specificity foundation model (SFM)^62–64^, as the binding verifier (**Extended Data Fig. 6a**). SFMs are thermodynamically-derived contrastive dual-encoder architectures that jointly embed ligand and receptor sequences to predict their interactions^62–64^. By explicitly modeling both interaction partners, a SFM trained on Fc:Fc-receptor interactions provides an orthogonal approach for aligning FcGPT with target Fc-profiles, complementing the traditional classifiers. Across both design strategies we generated more sequences for profiles expected to be particularly challenging to achieve. Combining both design pools and removing duplicates within each profile yielded 1,341,341 unique candidate designs. We then selected 20-200 Fc-variants per profile (**Fig. 6a**) for experimental validation, prioritizing predicted agreement with the target binding profile, predicted expression (**Extended Data Fig. 6b**), and sequence diversity.

**Fig. 6.**
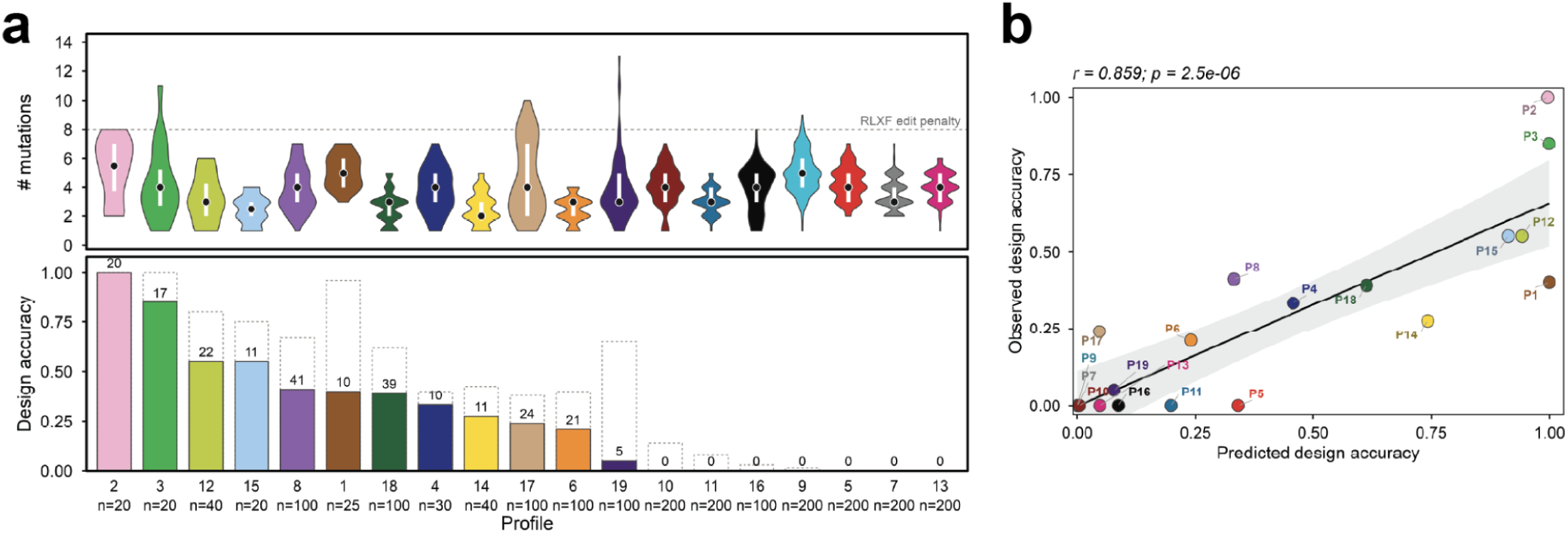
Experimental validation oí FcGPT’designtd Fc variants. a. Mutational load and experimental design accuracy across nineteen target Fc-receptor binding profiles, ordered by strict design accuracy. Top. Distribution of amino acid substitutions relative to the parental 299A-IYG sequence with median (dot) and interquartile range (white bar}. Dashed line indicates the threshold of eight substitutions beyond which the RLXF mutational penalty applies. Bottom, Design accuracy under strict (solid bars) and lenient (dashed outlines) scoring. Strict scoring requires all eight Fc-receptor binding labels to match the intended profile, whereas lenient scoring evaluates only receptors with definitive experimental labels* allowing sequences with fewer than eight resolved labels to count as correct (e g-* 7/7 or 6’6 matches), Numbers above bars indicate the number of successful designs under strict scoring; n indicates the total number of sequences tested per profile, b, Pearson correlation between predicted and experimentally observed design accuracy (Strict). Solid line indicates linear regression. with 95% shaded confidence interval in grey.

The final 2000 sequences were synthesized as part of a multiplexed gene fragment pool (Twist), cloned into yeast for surface display, and sorted via FACS to determine if they exhibited the designed Fc-receptor binding profiles (**Extended Data Fig. 6c**). Remarkably, deep sequencing of the sorted populations revealed that 100% of the designed variants were detected in the expression-positive population, despite typically harboring 2-6 amino acid substitutions relative to the parental 299A-IYG sequence (**Fig. 6a**). Some variants even exceeded 10 substitutions (**Fig. 6a**), demonstrating that FcGPT can generate highly mutated Fc sequences while preserving protein surface expression.

We evaluated whether the expressed variants achieved their intended binding outcomes by calculating the exact match, defined as the percent of sequences for which all experimentally determined Fc-receptor labels matched the design specification. By this stringent metric, FcGPT successfully designed variants for 12 of the 19 intended Fc-receptor binding profiles, with design accuracies ranging from 5% to 100% among successful objectives (**Fig. 6a**). For instance, the model achieved 100% design accuracy for FcγR1-selective binding (Profile 2) and 85% for the Fcγ receptor silenced Profile 3 (**Fig. 6a**).

Design accuracy varied across objectives, with selectivity between closely related Fc-receptors presenting particular challenges, consistent with our computational predictions (**Fig. 5e**). Indeed, predicted and experimentally observed design accuracies were strongly and significantly correlated (r = 0.859, p = 2.5×10^−6^) (**Fig. 6b**), demonstrating that FcGPT can predict *a priori* which binding profiles are easier or harder to design. Importantly, FcGPT was nevertheless able to achieve several of these challenging receptor specificities. The model successfully designed 24 variants that selectively bound FcγR2B and not FcγR2A (Profile 17) (**Fig. 6a**). It also achieved bidirectional selectivity between the highly homologous FcγR3A and FcγR3B receptors, designing 39 variants selective for FcγR3A (Profile 18) and five variants with the reciprocal FcγR3B-selective profile (Profile 19) (**Fig. 6a**). Together, these results demonstrate that FcGPT can generate proteins with desired patterns of binding activity, extending protein design from individual targets to entire families of receptors.

## DISCUSSION

This work unites high-throughput experimental screening and protein engineering with deep learning to establish a new paradigm for studying and engineering antibody biology. Traditional Fc engineering strategies have used experimental setups in which Fc libraries are generated and selected for Fc-receptor binding to isolate individual variants with desired phenotypes^12–14,16^ – a process that can take years. While effective for their intended goal, these approaches are inherently limited – they focus on identifying one or a few “hits” and discard the vast majority of valuable data generated during selection, including information about which variants bind, which do not bind, and the sequence features that explain why. By contrast, our deep learning models use large-scale experimental data to learn the sequence determinants of Fc-receptor recognition, then invert these relationships to design novel Fc-variants with desired binding profiles. Our experimental Fc library is 10^8^ in size, yet our models can access a theoretical sequence space of 10^62^ Fc-variants – tens of orders of magnitude beyond even the most high-throughput experimental techniques – unlocking *in silico* candidate prioritization, functional effect prediction, and hypothesis generation across a massive sequence space. We note, however, that the performance of our predictive and generative models may not generalize to the entire universe of Fc-variants, such as variants with mutations at positions not included in our library.

FcGPT decouples Fc engineering from the constraints of experimental biology. Instead of months to years of experimental screening, it enables the *in silico* generation of millions of Fc-variants with tailored, multi-receptor binding profiles. We challenged FcGPT by asking it to design Fc-variants across 19 distinct Fc-profiles – including some that have been notoriously difficult to engineer (e.g., FcγR2A/2B- and FcγR3A/3B-selectivity), and several of which we did not know were even biophysically possible from our starting Fc scaffold. Remarkably, despite having to achieve the desired binding pattern across all eight receptors at once, FcGPT successfully generated variants matching 12 of 19 target profiles, with design accuracies ranging from 5% to 100% among successful objectives – a success rate readily compatible with even low-throughput experimental validation. Although design accuracy varied across objectives, the strong correlation between predicted and observed design accuracy highlights that FcGPT can predict which objectives will be difficult *before* experimental testing. This emergent capability of the model will help prioritize experimental resources, guide the collection of additional training data, and ultimately reveal both the possibilities and constraints of Fc-receptor recognition in future work.

Beyond Fc biology, the general strategy underlying FcGPT brings a new capability to the field of protein design. Whereas leading structure-based methods, including RFdiffusion and BindCraft^65,66^, have primarily focused on designing proteins that bind individual targets, FcGPT tackles the considerably more complex challenge of simultaneously programming binding (and non-binding) across an entire family of receptors. This challenge is further compounded by the close similarity of the Fc-receptor system – a regime with nearly identical molecular targets. That FcGPT achieved experimental success rates comparable in magnitude to leading single-target design methods (e.g., 10-100% for BindCraft) is particularly notable given the added challenge of programming binding and non-binding across eight homologous receptors simultaneously. More fundamentally, our approach demonstrates that the desired “profile” of molecular recognition, rather than binding to an individual target, can itself become the objective of design. This new capability could be naturally extended beyond Fc-receptors to other families of immune proteins, opening new possibilities for encoding precise instructions for immune responses directly into engineered proteins.

## METHODS

### Yeast antibody Fc surface display cloning

Human IgG1 Fc sequences were ordered as custom, *S. cerevisiae* codon optimized gene fragments (Twist Bioscience) and cloned into the yeast surface display vector (pYD1) by Gibson assembly^67^. The pYD1 plasmid was modified such that the entire fusion to Aga2 was replaced with a cassette encoding a human IgG1 Fc domain, expression tags, and stop codon (HA tag-Fc-FLAG-Stop). Successful integration of the antibody Fc sequences was verified by Sanger sequencing.

### Antibody Fc deep mutational scanning library construction

Single-site deep mutational scanning (DMS) libraries of an aglycosylated human IgG1 Fc domain (299A-IYG)^48^, were generated using a nicking mutagenesis approach as described previously^68^. Mutagenesis was achieved using custom oligo pools ordered from Integrated DNA Technologies (IDT) containing degenerate NNK codons that tiled across each position of the 299A-IYG Fc domain spanning from residue 216 to 447 (EU numbering). The alanine at position 299 was kept constant to preserve the aglycosylated framework of the Fc region. The entire human IgG1 antibody Fc domain cannot be covered with a single Illumina sequencing read. Thus, two libraries – library 1 and library 2 – were individually constructed to cover the first and second half of the Fc domain respectively. The DMS libraries were constructed separately, but all subsequent steps were performed in a pooled manner. Following mutagenesis, the libraries were transformed into *E. coli* DH5-alpha ElectroMAX (ThermoFisher, 11319019) via electroporation for selection and expansion. Antibody Fc DMS library plasmid was extracted from *E. coli* according to the instructions of the manufacturer (Zymo, D4200). The antibody Fc DMS library plasmid was drop dialyzed for 2 hours using nuclease-free water (Invitrogen, AM9930), and transformed into *S. cerevisiae* yeast (EBY100; ATCC, MYA-4941) as previously described^69^. Following yeast library transformation, the cells were grown in SD-CAA medium [20g/L glucose (Sigma, G8270), 8.56 g/L NaH_2_PO_4_·H_2_O (Roth, K300.1), 6.77g/L Na_2_HPO_4_·2H_2_O (Sigma, 1.06580.0500), 6.7g/L yeast nitrogen base without amino acids (Sigma, Y0626) and 5g/L casamino acids (Gibco, 223120) in deionized water] for 2 days at 30°C with mild agitation.

### Screening antibody Fc libraries for Fc-receptor binding

Yeast cells containing antibody Fc library plasmid were cultured in SD-CAA medium for 18-24 hours at 30°C with mild agitation. To induce antibody Fc surface display, the cells were transferred into SG-CAA medium [20g/L galactose (Sigma, G0625), 8.56g/L NaH_2_PO_4_·H_2_O (Roth, K300.1), 6.77g/L Na_2_HPO_4_·2H_2_O (Sigma, 1.06580), 6.7g/L yeast nitrogen base without amino acids (Sigma, Y0626) and 5g/L casamino acids (Gibco, 223120) in deionized water] and incubated at 30°C with mild agitation for 24 hours. Subsequently, 5-10 million cells from the antibody Fc DMS libraries, or 1×10^7^ – 10×10^7^ cells from the antibody Fc combinatorial libraries, were centrifuged at 6000 x g for 3 minutes and washed twice in Wash Buffer [0.5% bovine serum albumin (Sigma, A2153), 2mM EDTA (Biosolve Chemicals, 05142391), and 0.1% Tween20 (Sigma, P9416) in PBS (Pan Biotech, P04-53500)]. Streptavidin-PE-Fc-receptor detection reagent was generated by mixing streptavidin-PE (Sigma, 42250) with a biotinylated Fc-receptor (ACROBiosystems: FcγR1 (FCA-H82E8), FcγR2A-H (CDA-H82E6), FcγR2A-R (CDA-H82E5), FcγR2B (CDB-H82E0), FcγR3A-F (CDA-H82E8), FcγR3A-V (CDA-H82E9), FcγR3B (CDB-H82E4), FcRn (FCM-H82W7)) in a 1:4 molar ratio. These were incubated for 10 minutes at room temperature (RT) in the dark shaking. To quench excess streptavidin, 500 μM d-biotin (Avidity) was added at a 1:10 ratio relative to the total volume. The yeast cells were then resuspended in the streptavidin-PE-Fc-receptor detection reagent (Concentrations: FcγR1 (1μg/mL), FcγR2A-H (1μg/mL), FcγR2A-R (1μg/mL), FcγR2B (1μg/mL), FcγR3A-F (5μg/mL), FcγR3A-V (5μg/mL), FcγR3B (20μg/mL), FcRn (10μg/mL)) and incubated shaking at RT for 1 hour in the dark. This was followed by two washes. Next, cells were treated with 1:100 diluted anti-FLAG AF647 (BioLegend, 637316), and incubated shaking at RT for 30 minutes in the dark. Following this incubation, the cells underwent two additional washes and were finally resuspended in Wash Buffer for fluorescence-activated cell sorting (FACS) on a BD FACSAria Fusion or BD FACSDiscover S8. FcRn-stained samples were stained in and washed at pH6. Populations of yeast cells were sorted based on their Fc-receptor binding ability, then centrifuged at 2000 x g for 5 minutes to remove the buffer. The cells were then resuspended in SD-CAA medium and grown for two days at 30°C with mild agitation. To maximize population purity, up to three successive sorting rounds were conducted for each Fc-receptor.

### Co-staining to screen for Fc-receptor selectivity

To enrich for Fc-variants with selective FcγR binding, the Fc combinatorial library was first enriched for FcRn binding at pH 6.0, followed by pairwise co-staining with six Fc-receptor combinations: FcγR1 × FcγR2A-R, FcγR1 × FcγR2B, FcγR1 × FcγR3A-F, FcγR2A-R × FcγR2B, FcγR2A-R × FcγR3A-F, and FcγR2B × FcγR3A-F. For each pair, biotinylated Fc-receptors were separately complexed with streptavidin-PE (Sigma, 42250) or streptavidin-APC (Biolegend, 405207) at a 1:4 molar ratio, using twice the receptor concentrations described for single-receptor staining. Following quenching with d-biotin, the two detection reagents were mixed at a 1:1 volume ratio immediately before staining, yielding the same final receptor concentrations as in single-receptor staining. Approximately 2×10^7^ induced yeast cells were resuspended in the combined detection reagents and incubated shaking at RT for 1 hour in the dark. This was followed by two washes. Next, cells were treated with 1:150 diluted anti-FLAG AF488 (BioLegend, 637318) and incubated shaking at RT for 30 minutes in the dark. Following this incubation, the cells underwent two additional washes and were resuspended in Wash Buffer for FACS. To enrich for receptor-selective Fc-variants, display-positive yeast cells were sorted from the two single-positive quadrants, corresponding to binding to one receptor but not the other. Sorted cells were resuspended in SD-CAA medium and grown for two days at 30°C with mild agitation. Multiple rounds of sorting were conducted to improve population purity. Purity was assessed by separately staining aliquots of the sorted populations with each receptor. Populations that remained impure after three rounds were further enriched by single-receptor sorts.

### Deep sequencing of antibody Fc DMS libraries

The antibody Fc plasmid library was extracted from yeast cells according to the instructions of the manufacturer (Zymo, D2004). Subsequently, two consecutive PCR reactions were performed to prepare the library for deep sequencing. The first PCR mixture contained NEBNext Ultra II Q5 Master Mix (New England Biolabs, M0544X), 50ng of library template DNA, and custom primers for library 1 (first half of the Fc domain) and library 2 (second half of the Fc domain) which possessed an 8 base pair unique molecular identifier (UMI) and Illumina partial adaptor sequences. PCR 1 conditions were: initial denaturation at 98°C for 5 minutes, followed by 5 cycles of denaturation at 98°C for 30 seconds, annealing at 64°C (Library 1) or 62°C (Library 2) for 90 seconds, extension at 72°C for 30 seconds, and a final extension at 72°C for 5 minutes. 2µL of Exonuclease I (New England Biolabs, M0293L) was added to each PCR product, followed by incubation at 37°C for 75 minutes to remove excess primers. Post-exonuclease, PCR products were cleaned-up using SPRIselect beads (Beckman Coulter, B23318) according to the instructions of the manufacturer. For the second PCR, the mixture contained NEBNext Ultra II Q5 Master Mix (New England Biolabs, M0544X), 10ng of library template DNA, and Nextera forward and reverse indexing primers (Illumina, 20027215). The second PCR amplification was similar to the first, but with an annealing temperature of 61°C and with 25 cycles. PCR products of the expected size were excised and gel extracted according to the instructions of the manufacturer (Zymo, D4002). Purified amplicons were pooled and assessed for sample quality using a Bioanalyzer, then subjected to paired-end sequencing using a MiSeq v3 Reagent Kit (600 cycles) (Illumina, MS-102-3003).

### Preprocessing of deep sequencing data

For the Fc DMS libraries, sequencing reads were paired, quality-trimmed, and merged in Geneious (v2023.0.4) using the BBTools suite with a quality threshold of Qphred R>20. Antibody Fc sequences were then extracted and processed using custom R (version 4.2.1) scripts. To correct for PCR amplification bias, sequences sharing identical unique molecular identifiers (UMIs) were collapsed into consensus sequences, which were then translated into amino acid sequences. Sequences containing N-linked glycosylation motifs (N-X-S/T) introduced during mutagenesis were filtered out. For the Fc combinatorial libraries, read pairing, trimming, and merging were performed using the same tools and quality parameters, but implemented via custom bash and Python (version 3.9.19) scripts to accommodate the larger file sizes.

### Deep mutational scanning analysis

Amino acid mutations in the deep sequencing data compared to the 299A-IYG wild type antibody were counted, and variants with zero or more than one mutation were removed. Variant count matrices, both pre- and post-sort, were then analyzed using R (version 4.2.1) for variant enrichment. Variants with fewer than five reads in the pre-sort (unselected) library were assigned a read count of 0 to reduce noise. A pseudocount of 1 was then added to each variant to prevent division by zero in subsequent steps. Pre- and post-selection matrices were normalized by their total read counts to account for differences in sequencing depth. Enrichment scores for individual variants were calculated by dividing the normalized post-selection frequency of each variant by its pre-selection frequency as described previously^70^. For positional enrichment scores, we specifically assessed the change in frequency of non-wild type amino acids at each position in the Fc domain. The normalized post-selection frequency of non-wild type amino acids at each position was divided by their pre-selection frequency, isolating the effect of selective pressures on the variation of each residue as previously described^70^. Fisher’s exact tests were used to assess statistical significance of enrichment at each position. A Bonferroni-adjusted p < 0.05 was considered statistically significant.

### Antibody Fc combinatorial library design

Results from the antibody Fc DMS screen were used to guide antibody Fc combinatorial library design. Specifically, residues that were significantly positively enriched in the non-binding population for each receptor – which are likely crucial for interaction with that receptor – were identified through positional enrichment analysis as described above. These residues were mutated using custom degenerate codons, tailored to reflect the amino acid distributions observed in the binding populations from the DMS data from each Fc-receptor. The selection of degenerate codons was based on minimizing the mean squared error between the degenerate codon, and the amino acid frequency observed in the binders, as described previously^23^. In instances where the degenerate codon could not encode the wild-type amino acid, frequencies were adjusted to ensure the inclusion of the wild type residue. Single-stranded DNA Ultramers incorporating degenerate codons at rationally-selected sites for targeted mutagenesis, and 299A-IYG (wild type) sequence at the remaining positions, were acquired from Integrated DNA Technologies (IDT). The combinatorial Fc library covered 137 amino acids of the Fc domain, spanning from residue 232 to 368 (EU numbering), and was partitioned into three segments to accommodate the length constraints of DNA Ultramers. The unassembled library, reflecting the diversity from eight different Fc-receptor DMS datasets, consisted of 24 mutant fragments and three 299A-IYG (wild type) segments. All fragments included BsmBI recognition sites for scarless Golden Gate assembly^55^, with standardized overhangs for each fragment type. Assembly fidelity was optimized using the NEB ligase fidelity viewer tool (https://ligasefidelity.neb.com/viewset/run.cgi).

### Antibody Fc combinatorial library construction

Prior to library assembly, a single-cycle PCR was performed to convert each DNA Ultramer fragment into double-stranded DNA, followed by gel extraction for purification. For Golden Gate assembly, the three 299A-IYG (wild-type) fragments were pooled to comprise 70% of the total insert pool (70% fragment 1, 70% fragment 2, 70% fragment 3), while the remaining 30% was evenly distributed among the eight mutational sublibrary fragments. Fragment inserts were combined with 2000ng of a custom entry vector (derived from pYTK090, Addgene #65197) at a 3:1 molar ratio (insert:vector). Using the NEBridge Golden Gate Assembly Kit BsmBI-v2 (New England Biolabs, E1602L), all library fragments were randomly assembled into full-length Fc gene segments in a single-pot Golden Gate digestion-ligation reaction^55^. To scale up the reaction, we used 26.67µL of Golden Gate Enzyme Mix and 53.34µL of T4 DNA Ligase Buffer. The assembly protocol consisted of 65 cycles of 42°C for 1 minute followed by 16°C for 1 minute, concluding with a final step at 60°C for 15 minutes. The assembled library was then transformed into *E. coli* MC1061F cells (Biosearch Technologies, 60514-1), resulting in 6×10^8^ transformants with a 93% rate of correctly assembled sequences. Plasmid DNA from the Fc combinatorial library was then extracted from *E. coli* cells according to the instructions of the manufacturer (Zymo, D4200). The assembled combinatorial Fc library was then PCR amplified, and a custom yeast display vector (based on pYD1) containing residues 369 to 417 of the Fc domain (EU numbering), was linearized using the restriction enzyme EcoRI (New England Biolabs, R3101L). Both the library insert and digested vector were gel purified according to the instructions of the manufacturer (Zymo, D4002). The Fc combinatorial library insert contained 81 base pair homology arms matching the linearized destination vector to enable assembly via homologous recombination in yeast. The library insert and linearized vector were co-transformed into *S. cerevisiae* yeast (EBY100; ATCC, MYA-4941) for assembly using a previously described protocol with minor changes^69^. In brief, EBY100 was grown overnight in YPD media [20g/L glucose (Sigma, G8270), 20g/L vegetable peptone (Sigma, 19942), and 10g/L yeast extract (Sigma, Y1625) in deionized water] at 30°C. The next day, cells from this culture were diluted into 500mL of YPD to reach an OD_600_ of 0.3 and grown until reaching an OD_600_ of approximately 1.6. 5mL of Tris-DTT [1M Tris pH8 (ThermoScientific, J22638.K2), 1M dithiothreitol (Sigma, D0632)] and 25mL of LiAc-TE [TE Buffer (Invitrogen, AM9849), 2M lithium acetate (Sigma, 517992)] were then added to the cells and incubated for another 15 minutes at 30°C with mild agitation to condition the cells, followed by three washes in cold Electroporation Buffer [0.6g Tris base (Sigma, T1503), 91.09g sorbitol (Sigma, S6021), 73.5mg CaCl_2_ dihydrate (Sigma, 1.02382.0500) in deionized water]. Electrocompetent EBY100 yeast cells were transformed with 83.33µg of the Fc library insert and 16.67µg of the linearized vector using 1mm electroporation cuvettes (Cell Projects, EP-101). Post-electroporation, cells were allowed to recover for 1 hour in YPD at 30°C with mild agitation, then transferred to selective SD-CAA medium for incubation overnight. After an overnight incubation at 30°C with mild agitation, the library concentration was adjusted back to an OD_600_ of 1 and subjected to another overnight growth phase. Colony-forming unit (CFU) plating was performed to assess library diversity, yielding 4.1×10^8^ transformants after three days.

### Deep sequencing of antibody Fc combinatorial libraries

The antibody Fc combinatorial plasmid library was extracted from yeast cells according to the instructions of the manufacturer (Zymo, D2004). Subsequently, two consecutive PCR reactions were performed to prepare the library for deep sequencing. The first PCR mixture contained NEBNext Ultra II Q5 Master Mix (New England Biolabs, M0544X), 80ng of template DNA, and custom designed primers which possessed an 8 base pair unique UMI, a 0-3bp heterogeneity spacer^71^, and Illumina partial adaptor sequences. PCR products were cleaned-up using SPRIselect beads (Beckman Coulter, B23318) according to the instructions of the manufacturer. For the second PCR, the mixture contained NEBNext Ultra II Q5 Master Mix (New England Biolabs, M0544X), 20ng of template DNA, and Nextera forward and reverse indexing primers (Illumina, 20027215). PCR products were cleaned-up using SPRIselect beads (Beckman Coulter, B23318). Purified amplicons were pooled and assessed for sample quality using a Bioanalyzer, then subjected to paired-end sequencing using a NovaSeq 6000 SP Reagent Kit v1.5 (500 cycles) (Illumina, 20028402) or a MiSeq v3 Reagent Kit (600 cycles) (Illumina, MS-102-3003).

### Antibody Fc structural analysis

Structural analyses were performed using PyMOL (Schrödinger, LLC). The human IgG1 Fc structure (PDB: 3AVE) was used as a reference^72^. Key structural regions on the antibody Fc domain were colored using a custom PyMOL (version 3.1.0) script. To visualize enrichment scores and mutational frequencies, values were extracted from a CSV file and mapped to individual Fc residues using a custom Python (version 3.9.19) script. The script scaled values to a color gradient and generated PyMOL commands for per-residue coloring. The resulting scripts were executed in the PyMOL (version 3.1.0) command line to display feature-specific residue patterns.

### Embedding Fc sequences using ESM2

We used the ESM2 protein language model (650M parameters)^32^, to generate embeddings of Fc sequences, extracting the CLS token representation as a fixed-length vector for each Fc-variant. PCA was then applied to the embeddings, and the first two principal components were visualized.

### Datasets and data splits

The training dataset comprised 1,474,696 unique Fc sequences collected across three experimental screens: (i) 324,092 sequences from deep mutational scanning (DMS), (ii) 1,101,665 additional sequences from the combinatorial library screen, and (iii) 48,939 additional sequences from the combinatorial co-staining screen. Each sequence was associated with up to eight binary Fc-receptor binding labels (binding or non-binding), with missing labels for receptors that were not experimentally evaluated. In total, 989,676 sequences had a single receptor-binding label, while 5,066 had labels for all eight receptors. Sequences were randomly partitioned into training (85%), validation (10%), and test (5%) sets, with the same partitioning strategy used for FcGPT pre-training. To further assess model generalization, we evaluated three more stringent splitting strategies: a combination-blocked split that held out entire combinations of mutated positions, a window-cluster-blocked split that held out mutation clusters across eight spatial windows of the Fc, and a position-blocked split that excluded 13 frequently mutated positions from training. Random splitting was retained for primary model benchmarking, while the additional splits were used to assess generalization to distinct mutational contexts (**Extended Data Table 2**). To further refine the models used for generative design, we experimentally characterized an additional 1,974 Fc variants generated using FcGPT via yeast display. These data were excluded from all model benchmarking analyses but were subsequently incorporated into the training of the Fc-receptor binding classifiers used as RLXF reward verifiers and for final sequence selection, as well as the expression predictor. The final FcGPT designs were generated following this refinement, and their experimental measurements were not used for model training or sequence selection.

### Fc-receptor binding classifiers

We trained an Fc-receptor-specific multi-layer perceptron (MLP) classifier to predict receptor-specific binding across each of the 8 Fc-receptors directly from the Fc amino acid sequence. The classifier had one shared trunk of weights across receptors. Sharing that trunk allowed the network to learn from the correlation between different receptors. The input is the 232-residue Fc sequence (E216-K447), one-hot encoded over the 20 canonical amino acids, flattened, and processed by a shared encoder of 4 hidden layers (*d_hid_* = 2, 048, ReLU activation, dropout p = 0.1), outputting eight logits - one per receptor -, that are passed through a sigmoid to yield the predicted binding probability for each receptor, *p_cls,r_* (Equation 1):

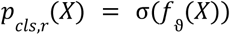

Equation 1. Output probability of binding for each Fc-receptor specific classification model; where:

- *p_cls,r_*(.) is the probability of a sequence to bind receptor r
- σ(.) is the sigmoid function
- *f*_ϑ_ (.) is the MLP whose *r*-th output is the logit for receptor *r*
- *X* = (*x_1_*, *x_2_*, *x_3_*, …, *x_r_*, …, *x_L_*) represents the amino acid sequence of the Fc region

Models were trained with an additive, receptor-wise binary cross-entropy (BCE) loss, l*_cls_*, that uses only the subset of labels present for each input (Equation 2):

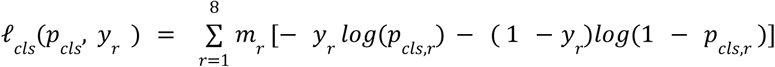

Equation 2. Additive binary cross entropy loss function *l_cls_* for the eight Fc-receptors used to train *p_cls_*; where:

- *y_r_* ɛ{0, 1, − 1} corresponding to true binding label for receptor r. −1 refers to no label.
- *m_r_* ɛ{0, 1} mask indicating wether *y* is known, returns 0 when *y* equals −1 and 1 otherwise

All model architectures were evaluated using identical held-out test sets. Models were trained using the Adam optimizer with a batch size of 128, early stopping with a patience of five epochs, and a maximum of 300 epochs. Hyperparameters were optimized using the tree-structured Parzen estimator (TPE) algorithm implemented in Optuna, with 200 trials of up to 50 epochs each, using the architecture with one output head per receptor. Model performance was evaluated across the random and three mutation-blocked data splits (**Extended Data Table 2**). Candidate reward verifiers were additionally evaluated on sequences with mutations at positions represented in the original library (in-zone) and at positions outside this space (off-zone) (**Extended Data Table 2**).

### FcGPT pre-training

To capture the distribution and high-order dependencies of the high-leverage positions in the 299A-IYG Fc domain, we trained FcGPT, a conditional autoregressive protein language model (PLM) pre-trained on Fc domain sequences. FcGPT is designed to model the conditional probability distributions associated with Fc-receptor binding profiles while simultaneously learning the amino acid distribution of the Fc region. The model takes as input amino acid sequences spanning residues E216 to K447 of the Fc domain, and prepends a fixed eight-token conditioning input prompt to each sequence. Each token in this prefix is a classification token (<yes>, <no>, or <pad>) representing the positive, negative and unknown binding outcome respectively for one of eight Fc-receptors (FcγR1, FcγR2A-H, FcγR2A-R, FcγR2B, FcγR3A-F, FcγR3A-V, FcγR3B, and FcRn). The order in which the tokens appear is critical, as the model associates each token position with a specific receptor and expects consistent positional encoding across both training and inference. FcGPT is trained to model the conditional residue distribution probability, *P_fc_* (Equation 3):

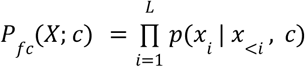

Equation 3. FcGPT model’s output probability of the next token given the Fc-receptor binding profile as an input; where:

- *L* denotes the sequence length
- *x*_r_ ɛ*A* represents that amino acid at position *i* drawn from the 20 standards amino acids Αɛ {*A*, *C*, *D*, *E*, *F*, *G*, *H*, *I*, *K*, *L*, *M*, *N*, *P*, *Q*, *R*, *S*, *T*, *V*, *W*, *Y*}
- *c* = (*c_1_*, *c_2_*, *c_3_*, *c_4_*, *c_5_*, *c_6_*, *c_7_*, *c_8_*) and *c_i_* ɛ {< *yes* >, < *no* >, < *pad* >} encodes the receptor-specific binding label
- *p*(.) conditional probability of amino acid *x_i_* given the input prompt and all the previous *x_<i_*
- *X* = (*x_1_*, *x_2_*, *x_3_*, …, *x_r_*, …, *x_L_*) represents the amino acid sequence of the Fc region

During training, the cross-entropy loss over the predicted amino acid vocabulary at each position is calculated and summed across the full sequence length to obtain the loss value, *P*, for each training step (Equation 4):

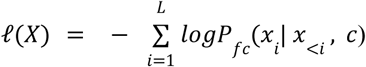

Equation 4. Cross-entropy loss function used during FcGPT pre-training; where:

- *x_r_* ɛ*A* represents that amino acid at position *r* drawn from the 20 standards amino acid
- *L* denotes the sequence length
- *X* = (*x_1_*, *x_2_*, *x_3_*, …, *x_r_*, …, *x_L_*) represents the amino acid sequence of the Fc region
- *P_fc_* (.) model’s output probability
- *c* = (*c_1_*, *c_2_*, *c_3_*, *c_4_*, *c_5_*, *c_6_*, *c_7_*, *c_8_*) and *c_i_* ɛ {< *yes* >, < *no* >, < *pad* >} encodes the receptor-specific binding label
- *x_r_* ɛ*A* represents that amino acid at position *i* drawn from the 20 standards amino acids Αɛ {*A*, *C*, *D*, *E*, *F*, *G*, *H*, *I*, *K*, *L*, *M*, *N*, *P*, *Q*, *R*, *S*, *T*, *V*, *W*, *Y*}

By encouraging FcGPT to accurately predict the next amino acid in the sequence given the input-condition prompt *c* and all previously observed residues *x*_<*r*_, the model captures the context-dependent distribution of Fc sequences and how it correlates with receptor binding profiles. Architecturally, FcGPT is a GPT-J based model following the ProGen2 framework^30^ (**Extended Data Fig. 5a**), and implemented as a 9,437,184-parameter sequence-only transformer. Each residue position is categorically encoded for all 232 residues, prepending the beginning of sequence (<bos>), an 8-token prompt corresponding to the binding profile of the variant and the end of sequence (<eos>) tokens, giving a total of *n*_pos_ = 242 input tokens. Rotary positional embeddings (*d_rot_* = 32) were applied to each token ^73^. The backbone consisted of a 3-layer transformer decoder block with an embedding dimension of *d_emb_* = 512 and *n*_ℎ*eads*_ = 32 self-attention heads, selected after evaluating multiple configurations and scales (**Extended Data Table 3**). A feed-forward network and softmax output layer predict the next token from a 32-character vocabulary. FcGPT uses causal (left-to-right) masking, enabling autoregressive sequence generation. The pre-training dataset comprised 1,474,696 unique sequences, split into 85/10/5 train/validation/test groups as for the classifiers. Model performance was evaluated based on validation cross-entropy loss and the edit distance between model-generated sequences and the wild type 299A-IYG (**Extended Data Fig. 5**). Scaling experiments across different parameter sizes showed that increasing model size beyond ∼10M parameters did not improve validation loss or edit-distance fidelity. Training was performed using the AdamW optimizer (β = 0.9,0.95; weight decay of 0.1), a learning rate of 3×10^−4^ with 500 warmup steps followed by cosined decay, gradient clipping at 1.0, bf16 precision, for 20,00 training steps, with early stopping upon convergence, a training batch size of 64, a validation batch size of 128. Specific architectural choices and configurations are shown in **Extended Data Table 3**. Ultimately, by predicting one amino acid at a time in a causally masked fashion conditioned on receptor prompts, FcGPT learns the sequence-function relationships of the Fc domain and enables generation of novel Fc sequences with user-specified Fc-receptor binding phenotypes.

### FcGPT post-training

To align FcGPT with high-accuracy Fc-receptor binding classifiers and enable the generation of Fc-variants with specific multi-receptor profiles, we developed a reinforcement learning-based prompt optimization pipeline. This approach was motivated by three factors: (i) the strong imbalance in binding profiles within the dataset, where some receptor profiles are well represented while others are rare or absent, (ii) the poor alignment between sequences generated by the pre-trained FcGPT base model and the classifier-predicted Fc-receptor binding profiles, and (iii) the prevalence of partial labels in the dataset (**Extended Data Fig. 4c**), which compounded sparsity and bias. Together, these factors skew the FcGPT base model *P_fc_* toward overrepresented profiles and reduce its capacity to capture rare but desirable or biologically important binding patterns. To overcome these limitations, we treated FcGPT as a policy model π*_fc_* and optimized it using an approach we term reinforcement learning with experimental feedback (RLXF), implemented with a Group Relative Policy Optimization (GRPO) framework ^37^. GRPO enables incorporation of a multi-reward function, guiding the policy model to generate sequences consistent with biological constraints and classifier expectations while regularizing divergence from the pre-trained distribution. GRPO further incorporates a KL divergence regularization term *D_KL_* between the fine-tuned policy π*_fc_* and the pre-trained model *P_fc_*, which serves to adapt the desired distribution while preserving the prior learned during pre-training. The objective function is (Equation 5):

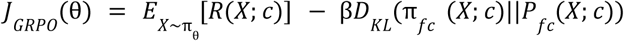

Equation 5. Loss function to be optimized for different binding profiles to guide the weight adjustment of the policy model, where:

- π*_fc_* (.) is the policy model
- *R*(.) is the reward function
- *D_KL_*(.) is the KL divergence function 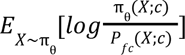
- β is a regularization term to balance the reward and the divergence term

Our reward function *R* (Equation 6) was constructed to: (i) promote sequences that match a specified Fc-receptor binding profile, (ii) constrain the mutational landscape by applying a threshold on edit distance from the wild type 299A-IYG variant to preserve therapeutic relevance while still allowing flexibility for the model to explore variants with potentially optimal binding properties, (iii) discourage reward exploitation by reducing credit for duplicate sequences within a batch, and (iv) penalize mutations at positions that the Fc combinatorial library did not vary a lot, which keeps designs where the classifiers are reliable.

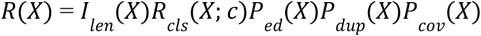

Equation 6. Multipurpose reward function used in GRPO to guide sequence generation for specific Fc-receptor binding profiles; where:

- *X* = (*x_1_*, *x_2_*, *x_3_*, …, *x_r_*, …, *x_L_*) represents the amino acid sequence of the Fc region
- *I_len_* (.) returns 1 if X equals the expected length, otherwise returns 0. Prevents policy model from exploring sequences with longer sequence length from the wild type.
- *R_cls_*(.) return a reward 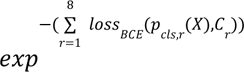 derived from classifier predicted probabilities. Higher agreement with the desired binding profile yields higher reward.
- *P_ed_*(.) returns 1 if X length is smaller than a threshold *ed_thr_*, otherwise returns 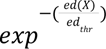. Penalizes sequences whose edit distance from the wild type exceeds a specific threshold.
- *P*_cov_(.) return a penalty 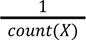. Reduces reward for sequences repeated within the same batch to prevent reward hacking from the policy model.
- *P*_cov_(.) returns *exp*^−2f^, where f is the fraction of mutations in X at positions outside the 42 positions varied in at least 1% of sequences in the library. Penalizes mutations at positions where classifiers are unreliable due to lack of data.

We sampled 100 completions per step and prompt, with a learning rate of 1 x 10^−4^, a KL coefficient of 0.04, 800 training steps, a sampling temperature of 0.8 and a maximum completion length of 234 tokens. While larger batch sizes would likely offer greater diversity by encouraging intragroup variability in the sequences generated, hardware constraints prevented further scaling. To adjust the weights of the FcGPT policy model during prompt optimization and to avoid catastrophic forgetting, we applied Low-Rank Adaptation (LoRA) for parameter-efficient fine-tuning ^59^. LoRA uses two low-rank matrices (*A* and *B*) in parallel with the transformer block to decompose the weight update matrix (Δ*W*) and reduce the number of trainable parameters (Equation 7):

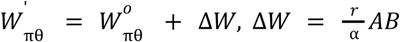

Equation 7. where:

- W’_πθ_ is the updated and W^0^_πθ_ is the pretrained πϑ weight matrices
- Δ*W* is the LoRA weight updates matrix
- *A*ɛℝ^dxr^ *and B*ɛℝ^rxd^ are the low rank matrices.
- *r* is the rank and α is the alpha
- *d* is the *d*_emb_ of the transformer block

The rank (*r*) controls the dimensionality of the low-rank update, and *alp*ℎ*a* (α) scales the update before it is added to the frozen pre-trained weights of the policy model. By updating only the newly introduced parameters while keeping the original pre-trained weights frozen, LoRA reduces memory and computational cost without causing catastrophic forgetting. We applied LoRA to the attention query, key, value, and output feed-forward modules of the policy model. We selected *r* = 32 after testing ranks of 16, 32, and 64; *r* = 16 decreased performance and *r* = 64 provided no additional benefit over 32. We used α = 64 as it represents a common choice in literature. Each specialist was trained on a single GPU with the TRL library for GRPO. The PLMFit package was used for parameter-efficient fine-tuning^74^, and the HuggingFace library for model management, training, and deployment^75^. This RLXF strategy produced prompt-specific LoRA adapters that improved conditional sequence generation, merging complementary information from FcGPT pre-training (inter-label correlations across receptors) and the receptor-specific classifiers (high-fidelity binding constraints). The result was a policy model capable of generating Fc sequences that both follow learned biological constraints and that align with classifier expectations. For each of the 19 target Fc-receptor binding profiles, we trained two RLXF specialists using distinct binding verifiers, yielding 38 specialists in total. The first strategy employed a coverage-gated ensemble of classifiers. For sequences with mutations predominantly at the 42 positions represented in at least 1% of the combinatorial library, binding predictions were obtained from the one-hot-encoded, shared-trunk MLP. For sequences extending beyond this well-represented mutational space, predictions were additionally evaluated using an ESM-C (300M) classifier trained with a supervised contrastive loss, taking the minimum predicted binding probability from the two models for each receptor. This conservative approach was intended to reduce overconfident binding predictions for sequences outside the training distribution. The second strategy employed the specificity foundation model (SFM), a contrastive dual-encoder built on frozen ESM-2 (650M) residue-level embeddings of Fc and receptor sequences^62–64^. The SFM was trained on 5,066 fully labeled sequences using a symmetric InfoNCE loss, with a two-parameter calibration mapping model compatibility scores to binding probabilities.

### Fc expression predictor

To prioritize FcGPT-generated sequences with favorable expression properties, we trained an expression predictor using 53,700 sequences from the combinatorial library and 33,500 sequences from the DMS library. Yeast displayed variants were separated into FLAG-positive and FLAG-negative populations by FACS, followed by deep sequencing. Sequences with fewer than three reads were excluded, and expression was quantified as the log2 enrichment between the two populations. We evaluated several model architectures and selected a multilayer perceptron (MLP) trained on frozen, mean-pooled ESM-C (300M) embeddings, which achieved comparable performance to the larger ESM-C (600M) model and outperformed alternative sequence-based expression scores (**Extended Data Fig. 6b**).

### Filtering and selection of FcGPT-generated sequences

For each of the 19 target Fc-receptor binding profiles, we generated between 20,000 and 500,000 sequences per RLXF specialist at a sampling temperature of 0.8, with larger sampling budgets allocated to more challenging profiles. Across both verifier strategies, this yielded 9.08 million candidate sequences, of which 1,341,341 remained after removing duplicates within each profile and strategy. We subsequently retained only sequences of wild type Fc length (232 amino acids) with mutations restricted to the variable region of the combinatorial library (EU residues 232-368), yielding 1,032,861 candidates. Each sequence was annotated with its predicted expression probability, mutational coverage within the training library, and presence in the experimental training dataset. Fc-receptor binding probabilities were evaluated using the shared-trunk, one-hot MLP classifier. Sequence selection was performed independently for each target profile and verifier strategy, with profile-specific quotas adjusted according to design difficulty. A small proportion of each quota was reserved for sequences with existing experimental binding measurements matching the target profile, while the majority was allocated to novel sequences absent from the training dataset. Novel candidates were prioritized based on predicted agreement with the target binding profile, first selecting exact matches across all eight receptors, followed by sequences matching seven of eight receptors. To promote sequence diversity, candidates were clustered into four groups based on their patterns of mutated positions using K means clustering, with sequences selected across clusters in a round-robin fashion. Within each cluster, preference was given to variants with mutations at well-represented library positions and higher predicted expression probabilities. Overlapping candidates between the two verifier strategies were assigned to only one strategy to avoid duplication. This process yielded a shortlist of 3,980 sequences, which was subsequently reduced to 2,000 sequences for experimental validation, comprising 1,995 FcGPT designs and five controls. The final designs included 186 sequences with existing experimental binding measurements, 1,232 novel sequences predicted to match all eight receptor-binding labels, 381 novel sequences matching seven of eight, and 196 additional candidates selected for challenging profiles.

### Validation of FcGPT-designed sequences

A validation set of 2000 FcGPT-designed Fc-variants was synthesized as part of a multiplexed gene fragment (MGF) pool (Twist Bioscience). Each Fc sequence contained 5′ and 3′ homology arms matching the linearized pYD1-derived yeast display vector to enable assembly via homologous recombination. The MGF pool was PCR-amplified using NEBNext Ultra II Q5 Master Mix (New England Biolabs, M0544X), 50ng of MGF template DNA, and custom primers. PCR conditions were: initial denaturation at 98°C for 30 seconds; 25 cycles of denaturation at 98°C for 20 seconds, annealing at 66°C for 20 seconds, and extension at 72°C for 30 seconds; followed by a final extension at 72°C for 5 minutes. The amplified pool was purified using a silica column clean-up kit (Zymo Research) and co-transformed with a custom linearized pYD1-derived yeast display vector into *S. cerevisiae* EBY100 for assembly by homologous recombination, as described above for the Fc combinatorial library, exceeding 1.0×10^8^ transformants. The resulting yeast-displayed FcGPT validation library was assayed for binding to the complete panel of Fc-receptors by FACS, and subjected to next-generation sequencing as described for the Fc combinatorial library. Fc-variants were classified as binders or non-binders based on at least fourfold enrichment in the binding or non-binding population, respectively.

### Hardware resources

All computational experiments were performed on Euler – the high-performance computing cluster at ETH Zurich. All models were trained and run on single NVIDIA RTX 4090 GPUs: FcGPT pre-training took about 3 h, classifier training up to 24 h and each RLXF specialist about 16 h, and generation and annotation ran as one GPU job per profile and arm.

## ACKNOWLEDGEMENTS

We gratefully acknowledge Thomas Horn, Mariangela Di Tacchio, Chiara Cavallini, Aleksandra Gumienny, and Svitlana Malysheva from the ETH Zürich Department of Biosystems Science and Engineering (D-BSSE) Single Cell Facility for their support with FACS screening. We additionally thank Christian Beisel, Mirjam Feldkamp, Elodie Burcklen, and Ina Nissen at the ETH Zürich D-BSSE Genomics Facility Basel for their outstanding assistance with next-generation sequencing.

## AUTHOR CONTRIBUTIONS

Conceptualization: E.B.I, W.K., S.T.R. Methodology: E.B.I., E.S., T.B., L.F., D.C., H.Y., C.R., M.M., G.B., D.M.M. Software: E.B.I., E.S., T.B., N.S., C.R. Validation: E.B.I., E.S., T.B., J.K., N.S., A.T. Formal analysis: E.B.I., E.S., T.B., A.K.W., N.S. Investigation: E.B.I., E.S., T.B., J.K., L.F., A.T., W.K., A.K.W., H.S., D.C., N.N., R.P., J.B., H.Y., M.M., G.B. Resources: N.N., R.P., J.B., S.T.R. Data curation: E.B.I, E.S., T.B., N.S. Writing – Original Draft: E.B.I. Writing – Review & Editing: all authors. Visualization: E.B.I., E.S., T.B., H.S. Supervision: E.B.I., W.K., S.T.R. Project administration: E.B.I, W.K., S.T.R. Funding: E.B.I, D.M.M., W.K., S.T.R.

## DATA AVAILABILITY

Data supporting the findings of this study are provided in the Article and Supplementary Information. The deep mutational scanning dataset, providing a high-resolution map of how amino acid substitutions across the antibody Fc domain affect binding to eight Fc receptors, will be made publicly available upon publication. The complete FcGPT training dataset is not publicly available due to intellectual property and commercialization considerations.

## CODE AVAILABILITY

The code used to develop the Fc-receptor binding classifiers and FcGPT is available on GitHub: https://github.com/LSSI-ETH/fc_engineering.

## EXTENDED DATA FIGURES & TABLES

**Extended Data Fig. 1.**
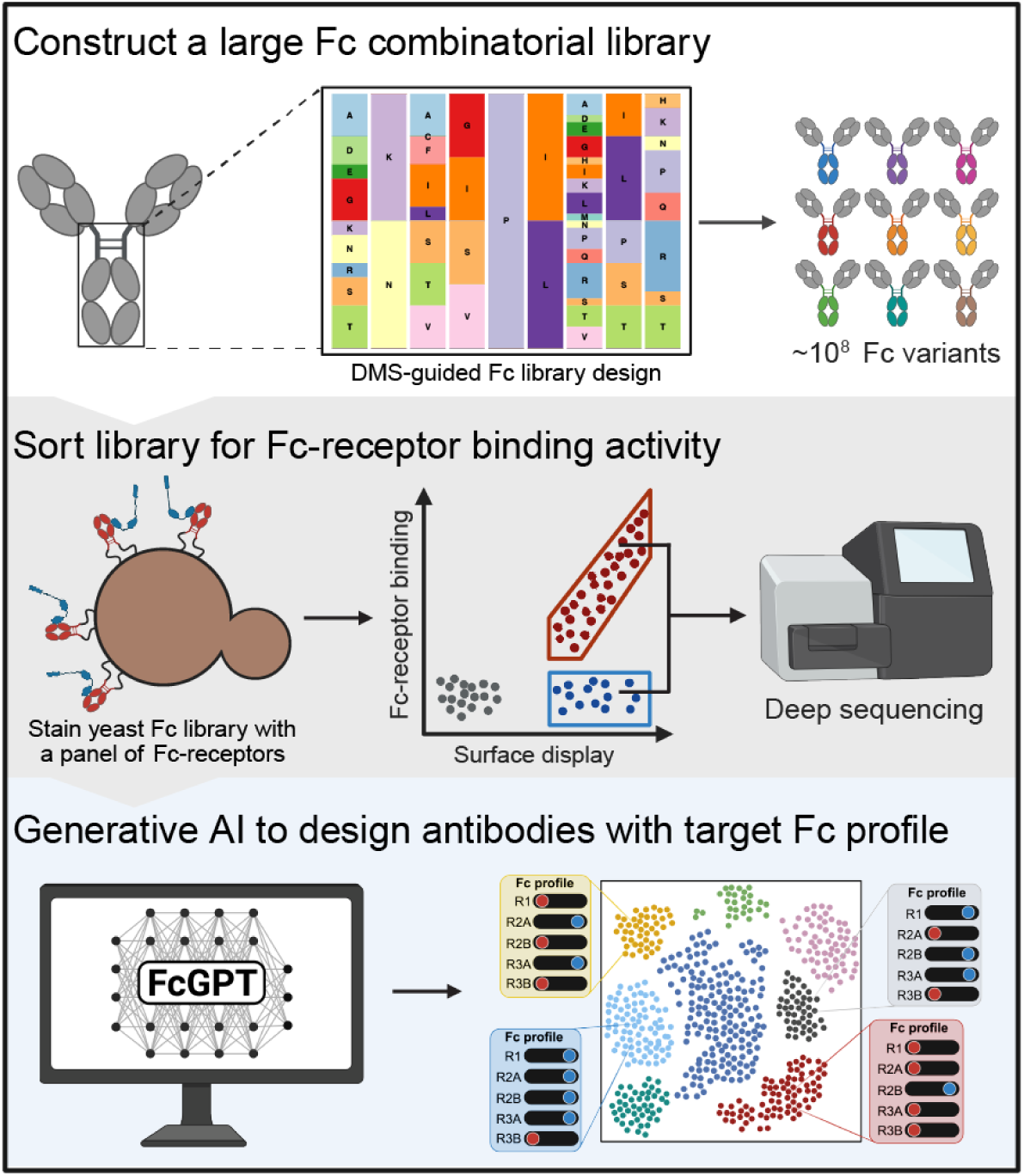
Developing a machine learning-guided platform for Fc engineering. Deep mutational scanning across the antibody Fc domain guided the construction of a large Fc combinatorial library (top). The library was screened for binding across a panel of Fc-receptors by yeast surface display, isolating binding and non-binding populations that were then deep sequenced (middle). These data were used to train FcGPT, a domain-specific protein language model able to computationally design antibodies with customized, user-defined Fc-receptor binding profiles (bottom).

**Extended Data Fig. 2.**
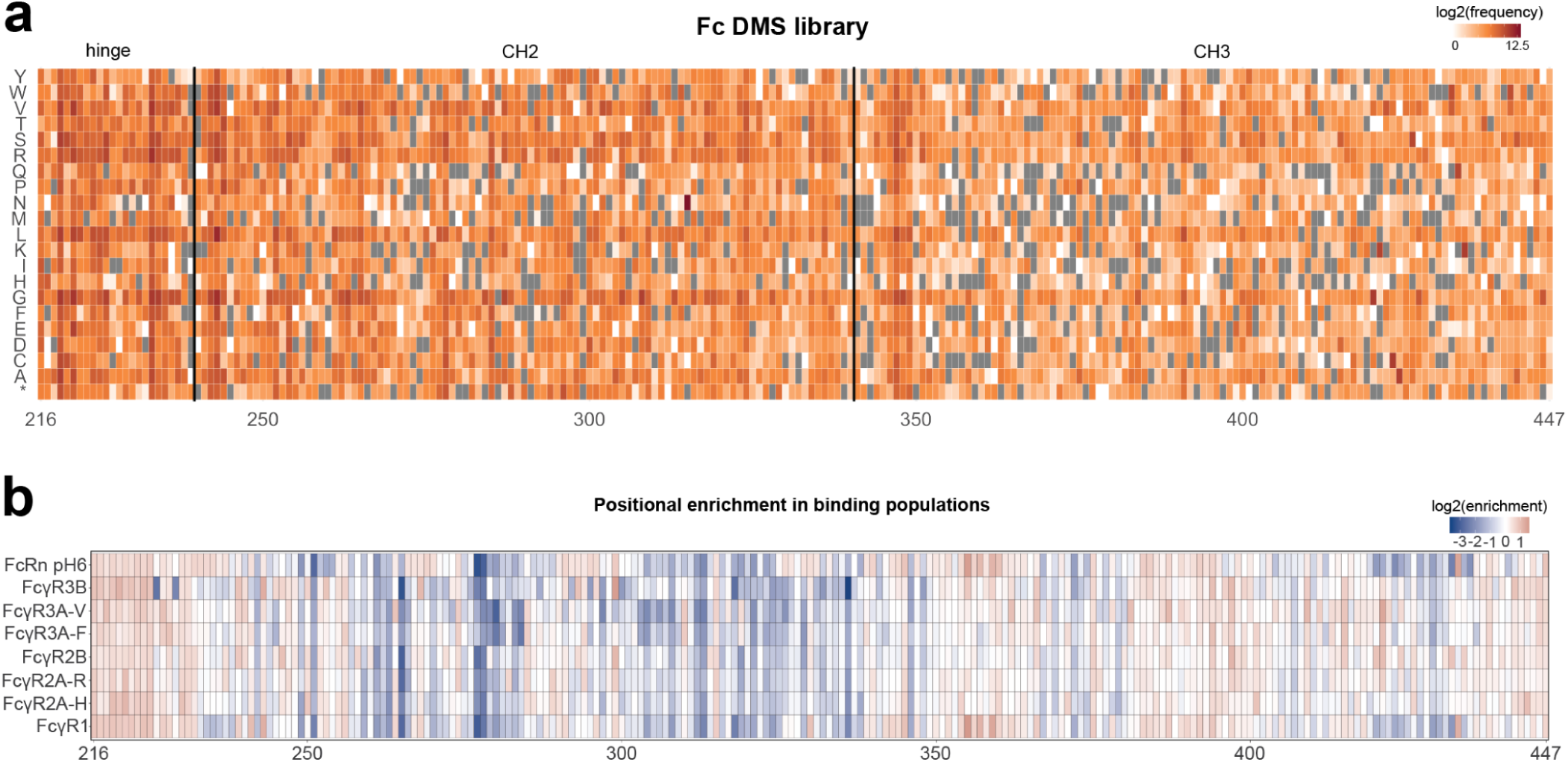
Deep mutational scanning library and enrichment. a, Heatmap depicting log2 frequency of mutations in the naïve yeast Fc DMS library. Rows correspond to amino acids, columns to Fc residues (EU numbering). **b,** Heatmap depicting log2 enrichment of mutations at each Fc position in the binding populations relative to the naïve library for eight Fc-receptors. Rows correspond to Fc-receptors, columns to Fc residues.

**Extended Data Fig. 3.**
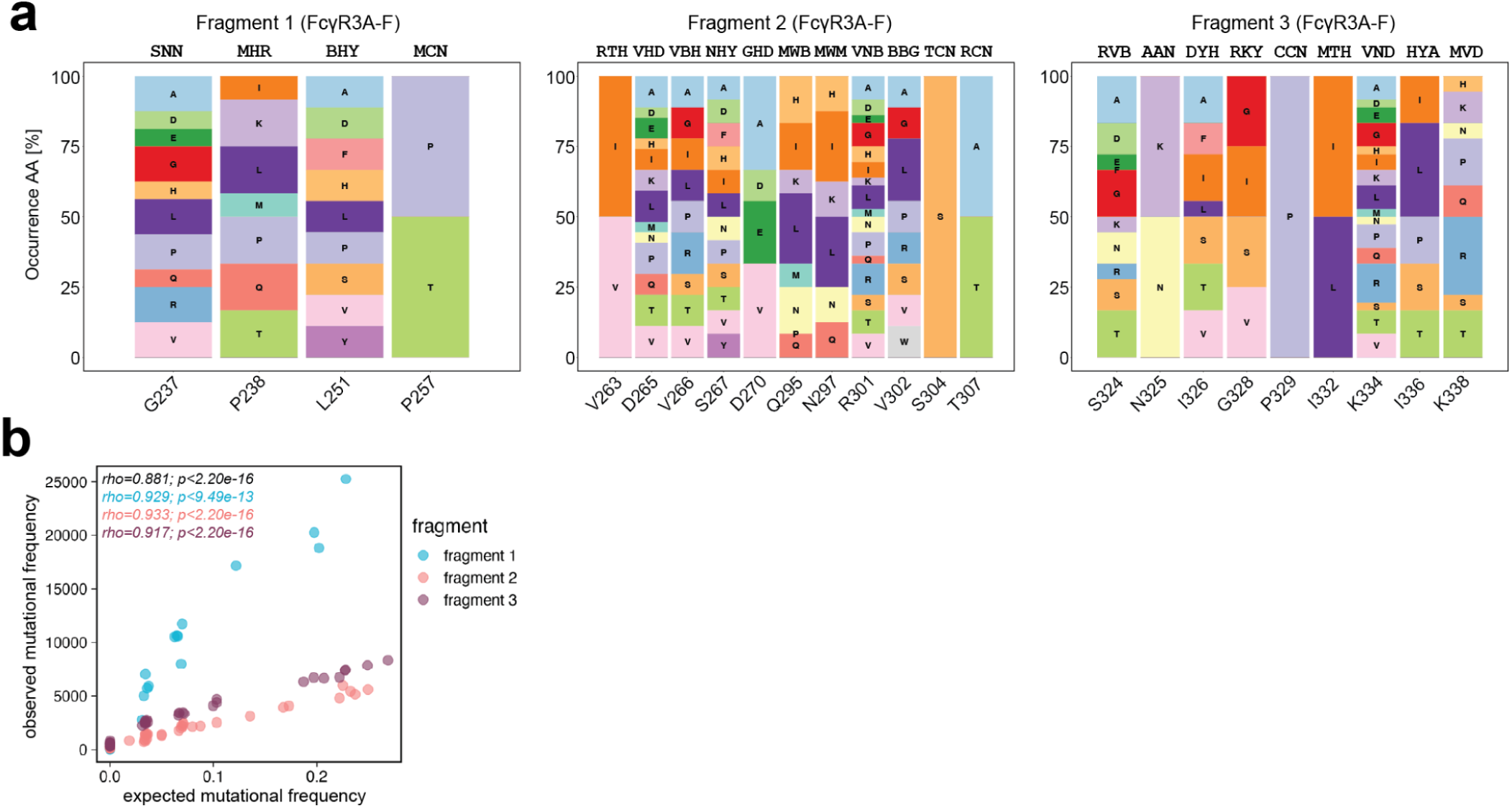
DMS-guided codon design and validation of the Fc combinatorial library. a, Amino acid distributions encoded by degenerate codons at selected positions across the three FcγR3A-F library fragments, shown as a representative example. Codons were selected based on amino acid frequencies in DMS binding populations. Positions not shown remained wild type. **b,** Spearman correlation (rho) and p-values are shown for each library fragment (colored) and overall (black), comparing the actual mutation frequency at each Fc amino acid position to the expected frequency based on the library design.

**Extended Data Fig. 4.**
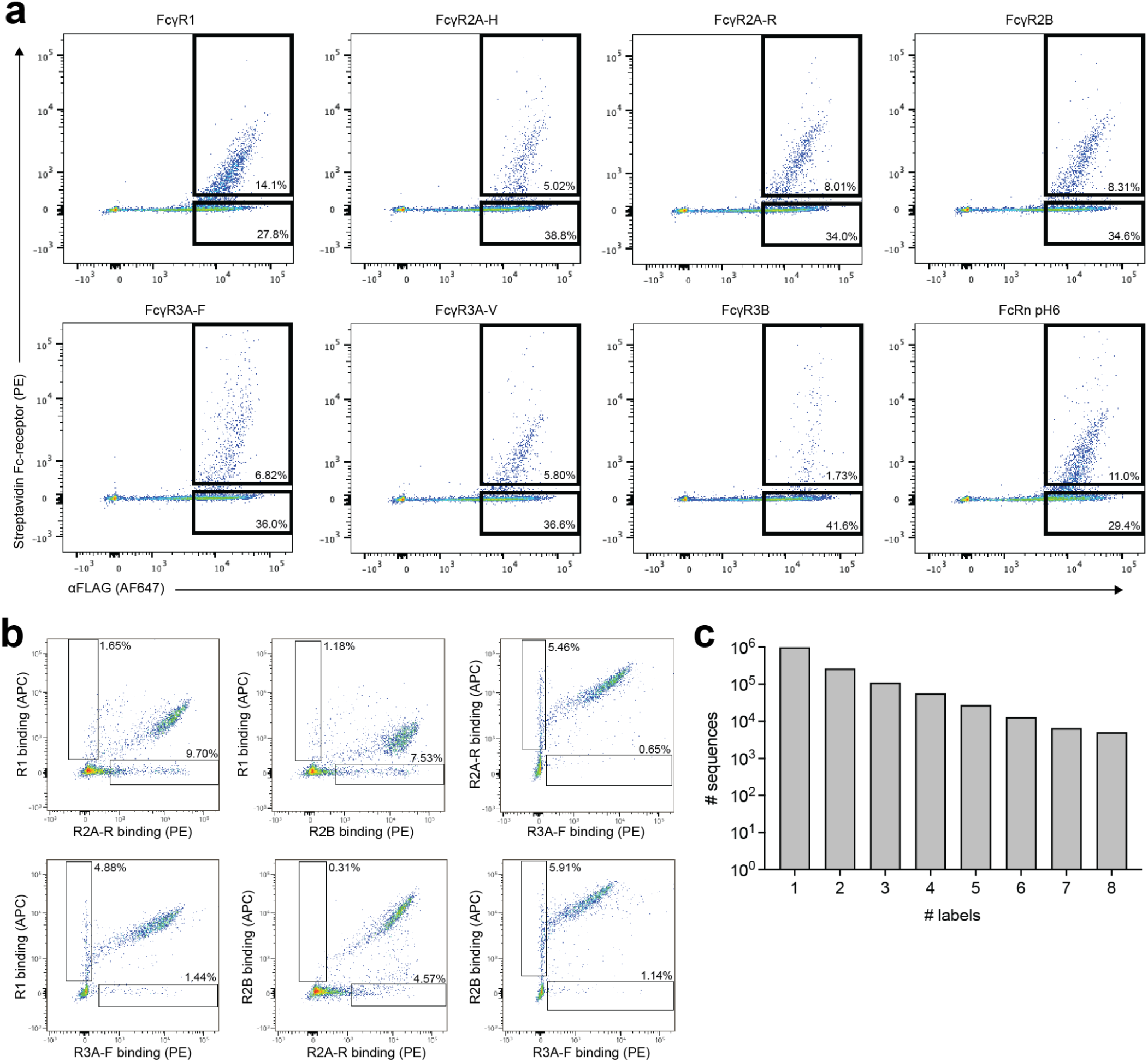
FACS and deep sequencing of the Fc combinatorial library. a, Flow cytometry plots showing Fc-receptor binding (streptavidin-PE) versus surface expression (αFLAG-AF647) during the first round of sorting. Gates indicate binding and non-binding populations, with percentages of cells indicated. Receptor concentrations were 1 μg/mL (FcγR1, FcγR2A-H, FcγR2A-R, FcγR2B), 5 μg/mL (FcγR3A-F, FcγR3A-V), 20 μg/mL (FcγR3B), and 10 μg/mL (FcRn, pH 6.0). **b,** Flow cytometry plots of pairwise Fc-receptor co-staining during the first round of sorting, with gates indicating receptor-selective populations and their frequencies. **c,** Distribution of unique Fc sequences by the number of receptors with experimentally determined binding labels.

**Extended Data Fig. 5.**
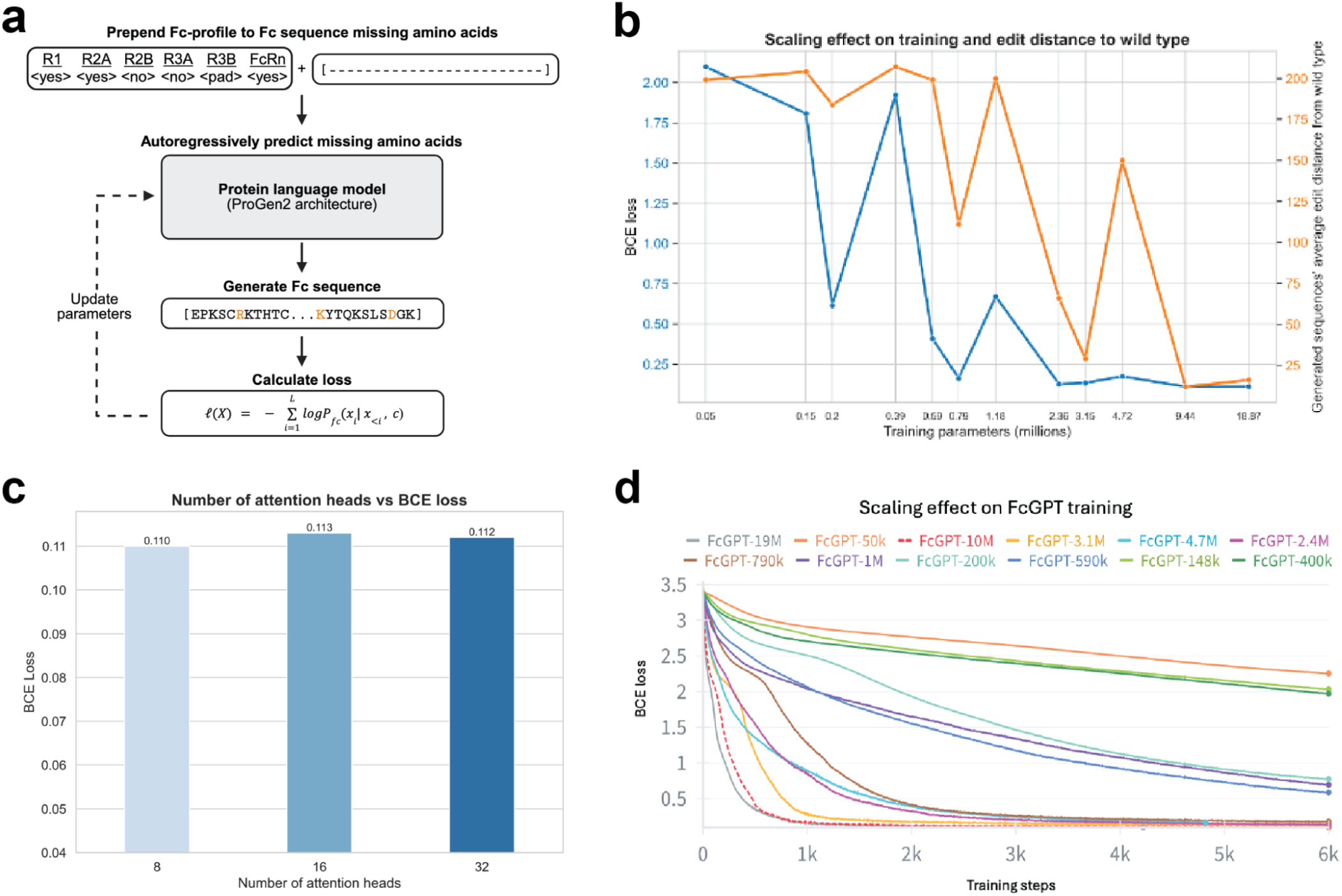
FcGPT pre-training. a, Each Fc sequence is prepended with an eight-token conditioning prompt encoding Fc-receptor binding outcomes (<yes>, <no>, or <pad>) for FcγR1, FcγR2A-H, FcγR2A-R, FcγR2B, FcγR3A-F, FcγR3A-V, FcγR3B, and FcRn (pH 6). The model is a 3-layer transformer with 32 self-attention heads, an embedding dimension of 512, and feed-forward layers followed by a softmax output that autoregressively predicts missing amino acids to generate candidate Fc sequences. Sequences are evaluated under a cross-entropy loss, and parameters are updated via gradient descent. See Methods for additional details. Image created with BioRender.com. **b,** Scaling effects on model performance, showing the relationship between model size (number of training parameters) with both the validation cross-entropy loss (left axis) and the average edit distance from the wild-type sequence for generated variants (right axis). **c,** Effect of different number of attention heads (8, 16, 32) during training shows minimal impact on validation cross-entropy loss. **d,** Training loss trajectories during FcGPT pre-training for model size scaling from 50k to 19M parameters. Larger models converge faster and achieve lower cross-entropy losses. Scaling beyond 10M parameters provides no additional benefit, leading to the selection of FcGPT-10M as the primary architecture.

**Extended Data Fig. 6.**
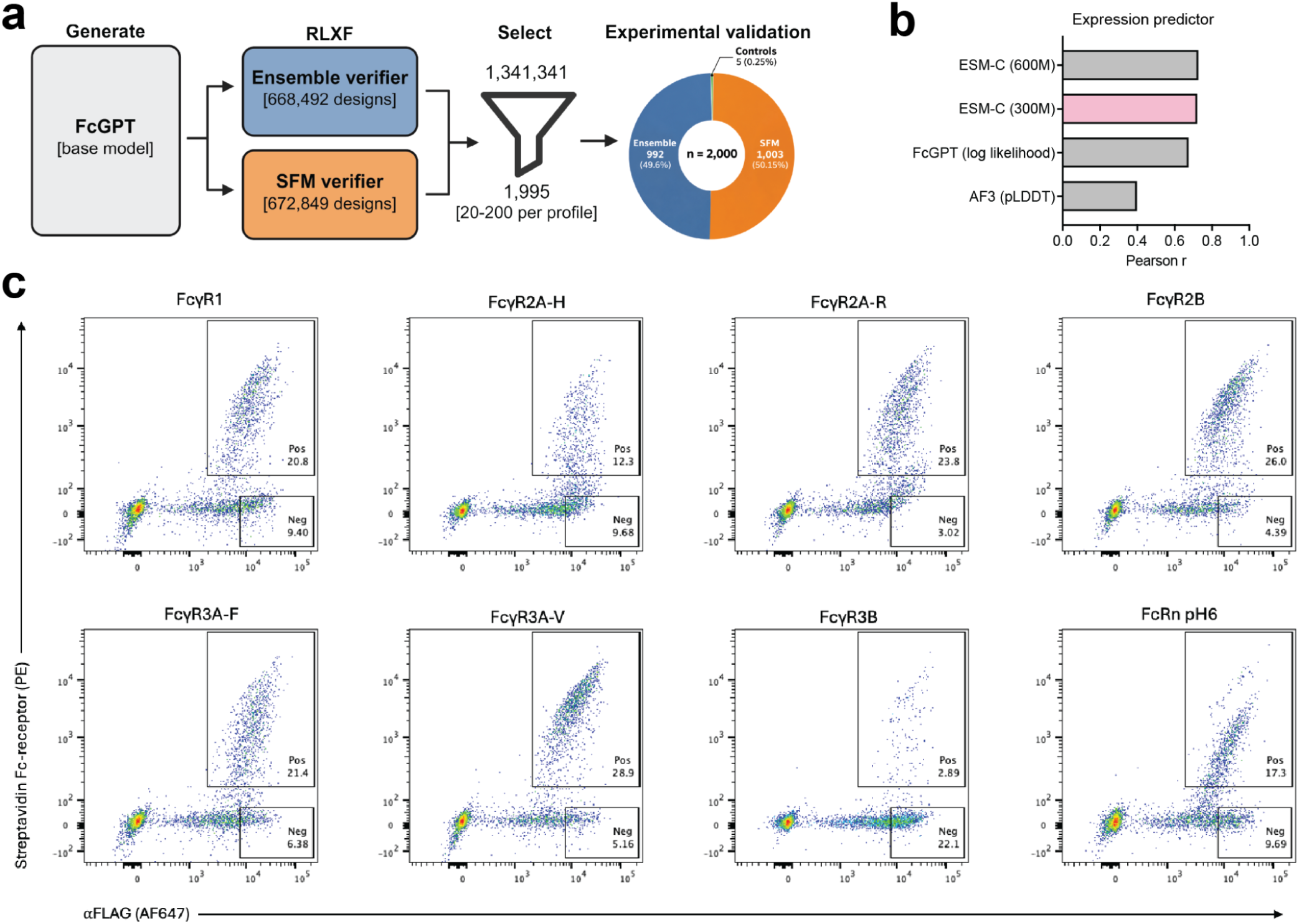
Generation, selection, and characterization of FcGPT-designed Fc variants. a, FcGPT design workflow using two parallel RLXF strategies with either an ensemble of Fc-receptor binding classifiers or a specificity foundation model (SFM) as the binding verifier. The two strategies yielded 1,341,341 unique candidate designs, from which 1,995 were selected for experimental validation alongside five controls. **b,** Pearson correlation between predicted and experimentally measured log₂ expression enrichment for 1,285 FcGPT-designed variants excluded from expression predictor training. Predictions were based on MLPs trained on frozen, mean-pooled ESM-C 300M or 600M embeddings, FcGPT sequence log-likelihood, or AlphaFold3 mean pLDDT. The ESM-C 300M model (pink) was selected for expression prediction. **c,** Flow cytometry analysis of the FcGPT-designed validation library during the first round of FACS sorting. Fc expression was measured using αFLAG-AF647 (x-axis) and Fc-receptor binding using streptavidin-PE (y-axis). Fc-receptors were used at the following concentrations: FcγR1 (1μg/mL), FcγR2A-H (1μg/mL), FcγR2A-R (1μg/mL), FcγR2B (1μg/mL), FcγR3A-F (5μg/mL), FcγR3A-V (5μg/mL), FcγR3B (20μg/mL), and FcRn (10μg/mL). FcRn staining was performed at pH 6.0. Gates indicate receptor-binding (Pos) and nonbinding (Neg) populations, with percentages shown.

**Extended Data Table 1.** Fc-receptor panel.

| Fc-receptor | Contact location | Cell distribution | Function(s) |
|---|---|---|---|
| FcγR1 | Lower hinge | Macrophages, monocytes, dendritic cells*, neutrophils* | Phagocytosis, antigen presentation, phagocyte activation |
| FcγR2A (167R)<br>low affinity allele | Lower hinge | Macrophages, monocytes, dendritic cells, neutrophils, eosinophils, basophils, mast cells | Phagocytosis, immune activation |
| FcγR2A (167H)<br>high affinity allele | Lower hinge | Macrophages, monocytes, dendritic cells, neutrophils, eosinophils, basophils, mast cells | Phagocytosis; immune activation |
| FcγR2B | Lower hinge | B cells, macrophages, monocytes, dendritic cells, eosinophils, basophils, mast cells | Limits immune activation |
| FcγR3A (176F)<br>low affinity allele | Lower hinge | Macrophages, monocytes, dendritic cells, NK cells | ADCC |
| FcγR3A (176V)<br>high affinity allele | Lower hinge | Macrophages, monocytes, dendritic cells, NK cells | ADCC |
| FcγR3B | Lower hinge | Neutrophils | Unknown |
| FcRn | CH2-CH3 junction | B cells, macrophages, monocytes, dendritic cells, neutrophils | Protects IgG from degradation, transcytosis |

\* *Indicates expression is inducible. <sup>2,3</sup>*

**Extended Data Table 2.**
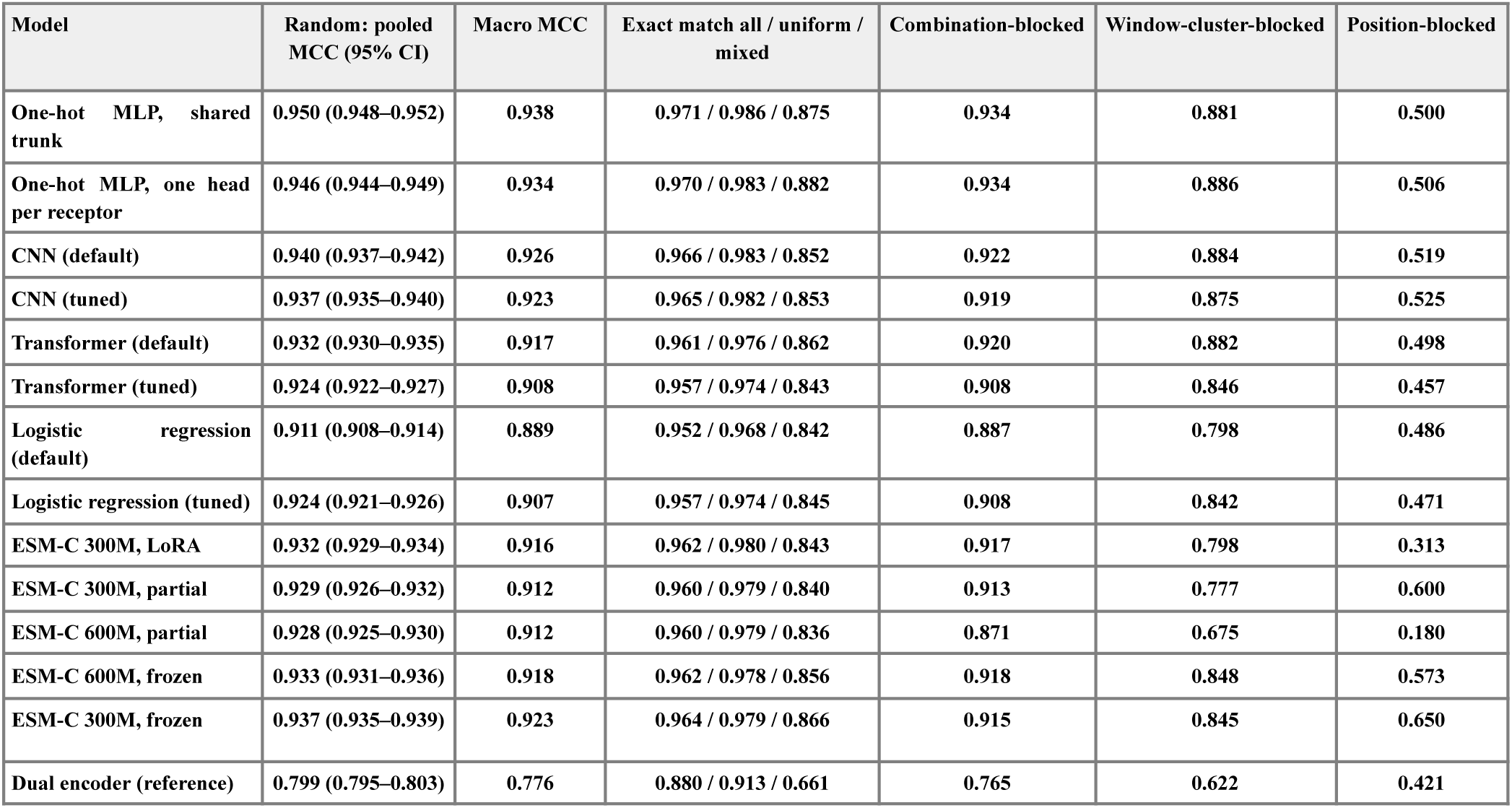
Performance comparison of different Fc-receptor classifier architectures. Pooled Matthews correlation coefficient (MCC, threshold 0.5) of each model on the held-out test set of four data splits; for the random split, 95% bootstrap confidence intervals, macro MCC (mean of the eight per-receptor MCCs) and exact match, the fraction of sequences for which all labeled receptors are predicted correctly (all sequences / sequences with uniform labels / sequences with mixed labels). One-hot MLPs, logistic regression and the CNN use flattened or one-hot sequences and the transformer token embeddings; ESM-C models use mean-pooled embeddings with an MLP head, with the encoder frozen, partially fine-tuned (last four blocks) or LoRA-fine-tuned. “Default” denotes fixed hyperparameters and “tuned” a 60-trial TPE search (200 trials for the one-hot MLPs). The dual encoder, trained only on 2.4–4.3k fully labeled sequences, is shown for reference.

**Extended Data Table 3.** Architectures cross-entropy loss after convergence during hyperparameter tuning for the selection of the final FcGPT configuration.

| Version | Embedding dimension | No. layers | No. parameters | BCE loss |
|---|---|---|---|---|
| FcGPT-50k | 64 | 1 | 49152 | 2,098 |
| FcGPT-148k | 64 | 3 | 147456 | 1,808 |
| FcGPT-400k | 64 | 8 | 393216 | 1,922 |
| FcGPT-200k | 128 | 1 | 196608 | 0,615 |
| FcGPT-590k | 128 | 3 | 589824 | 0,408 |
| FcGPT-1M | 128 | 6 | 1179648 | 0,67 |
| FcGPT-790k | 256 | 1 | 786432 | 0,162 |
| FcGPT-2.4M | 256 | 3 | 2359296 | 0,126 |
| FcGPT-4.7M | 256 | 6 | 4718592 | 0,173 |
| FcGPT-3.1M | 512 | 1 | 3145728 | 0,133 |
| FcGPT-10M | 512 | 3 | 9437184 | **0,11** |
| FcGPT-19M | 512 | 6 | 18874368 | **0,11** |

